# The kinase activity of the cancer stem cell marker DCLK1 drives gastric cancer progression by reprogramming the stromal tumor landscape

**DOI:** 10.1101/2022.04.21.489109

**Authors:** Shoukat Afshar-Sterle, Annalisa L E Carli, Ryan O’Keefe, Janson Tse, Stefanie Fischer, Alexander I Azimpour, David Baloyan, Lena Elias, Pathum Thilakasiri, Onisha Patel, Fleur M Ferguson, Moritz F Eissmann, Ashwini L Chand, Nathanael S Gray, Rita Busuttil, Alex Boussioutas, Isabelle S Lucet, Matthias Ernst, Michael Buchert

## Abstract

Gastric cancer (GC) is the 3rd leading cause of cancer mortality worldwide, therefore providing novel diagnostic and treatment options is crucial for at risk groups. The serine/threonine kinase doublecortin-like kinase 1 (DCLK1) is a proposed driver of GC with frequent amplification and somatic missense mutations yet the molecular mechanism how DCLK1 mediates tumorigenesis is poorly understood. We report how DCLK1 expression orchestrates complementary cancer cell intrinsic and extrinsic processes leading to a comprehensive pro-invasive and pro-metastatic reprogramming of cancer cells and tumor stroma in a DCLK1 kinase-dependent manner. Mechanistically, we identify the chemokine CXCL12 as a key promoter of the pro-tumorigenic properties downstream of DCLK1. Importantly, inhibition of the DCLK1 kinase domain reverses the pro-tumorigenic and pro-metastatic phenotype. Together, this study establishes DCLK1 as a promising, targetable master regulator of GC.

**Teaser:** DCLK1 is a druggable cancer driver of GC

## Introduction

More than one million new cases of gastric cancer (GC) occurred in 2018, making it the fifth most common malignancy globally and the world’s third leading cause of cancer mortality with almost 800,000 deaths in 2018 (*1*). Genomic profiling has identified 4 molecular subtypes of GC, chromosomal instable (CIN), microsatellite instable (MSI), Epstein Barr Virus-transformed (EBV) and genomic stable (GS) (*2*). Among the different subtypes of GC, GS have the highest stroma-to-tumor ratio, indicating a prominent desmoplastic response. Herein, a complex network of paracrine and autocrine signaling crosstalk between cancer cells and non-cancer cells shapes the tumor microenvironment (TME) and guides tumor evolution. In contrast, gastric tumors with stromal-low anatomy are mostly characterized by the activation of cell intrinsic oncogenic pathways that drive tumor neoplasia through a sequence of metaplastic and dysplastic events (*3*). The stroma-to-tumor ratio is clinically relevant in particular in GC where previous studies have shown that a low tumor-stroma ratio (stroma rich) is an independent prognostic factor in both intestinal and diffuse histological subtypes of GC (*4*).

Doublecortin-like kinase 1 (DCLK1) is a microtubule-associated serine/threonine kinase with tubulin polymerization activity (*5–11*). It is a putative cancer stem cell marker, a proposed prognostic marker for malignant GI tumors and a presumptive cancer driver of GC (*12–15*). DCLK1 promotes an epithelial to mesenchymal transition (EMT) in gastrointestinal (GI) cancer cells, driving disruption of cell-cell-adhesion, migration and invasion (*16–18*). Its EMT-promoting role in human cancer cells explains the strong link of DCLK1 as a marker of (cancer) stem cells or tumor initiating cells, particularly in cancers along the rodent GI tract (*16, 17, 19*). However, in these rodent models, Dclk1 is strongly expressed in epithelial tuft cells. Contrary to rodents, in humans, DCLK1 is not a marker for tuft cells as demonstrated by recent single cell studies (*20, 21*). Moreover, accumulating evidence suggests that DCLK1 may affect tumorigenesis in a cancer cell extrinsic manner through DCLK1’s ability to promote extracellular vesicle biogenesis emphasizing the strong correlation between DCLK1 expression and an immune suppressive tumor microenvironment in GI tumors (*22, 23*).

While our collective knowledge supports a model whereby DCLK1 promotes tumorigenesis through cell intrinsic and extrinsic pathways, the underpinning molecular mechanisms remain largely unknown. Here, we use both *in vivo* and *in vitro* models to map the functional contributions of DCLK1 to gastric cancer development and identify corresponding therapeutic vulnerabilities.

## Results

### DCLK1 expression pattern in human gastric cancer

In order to investigate the expression profile of DCLK1 in gastric cancer, we first interrogated the stomach adenocarcinoma dataset (STAD) of The Cancer Genome Atlas (TCGA). DCLK1 expression is significantly upregulated in gastric cancers belonging to the genomic stable (GS) molecular subtype, Lauren’s histological diffuse subtype and the immune-enriched, fibrotic (IE/F) tumor microenvironment subtype (*24*) (**Figure 1 A-C**). There is also a significant trend towards higher expression levels of DCLK1 in more aggressive cancers irrespective of whether cancers are classified according to TNM stage, neoplasm histology grade (G1-G3) or the AJCC’s invasiveness of primary tumor classification (**Figure 1 D-F**). In line with this trend, high DCLK1 expression correlated with poorer overall (OS), progression-free (PFS) and disease-specific survival (DSS) (**Figure 1 G-I**). Meanwhile, the proportion of DCLK1-high tumors among newly diagnosed gastric cancers remained mostly constant throughout the four decades (40y to 80y) covering the vast majority of GC cases (**Figure 1 J**).

**Fig. 1.**
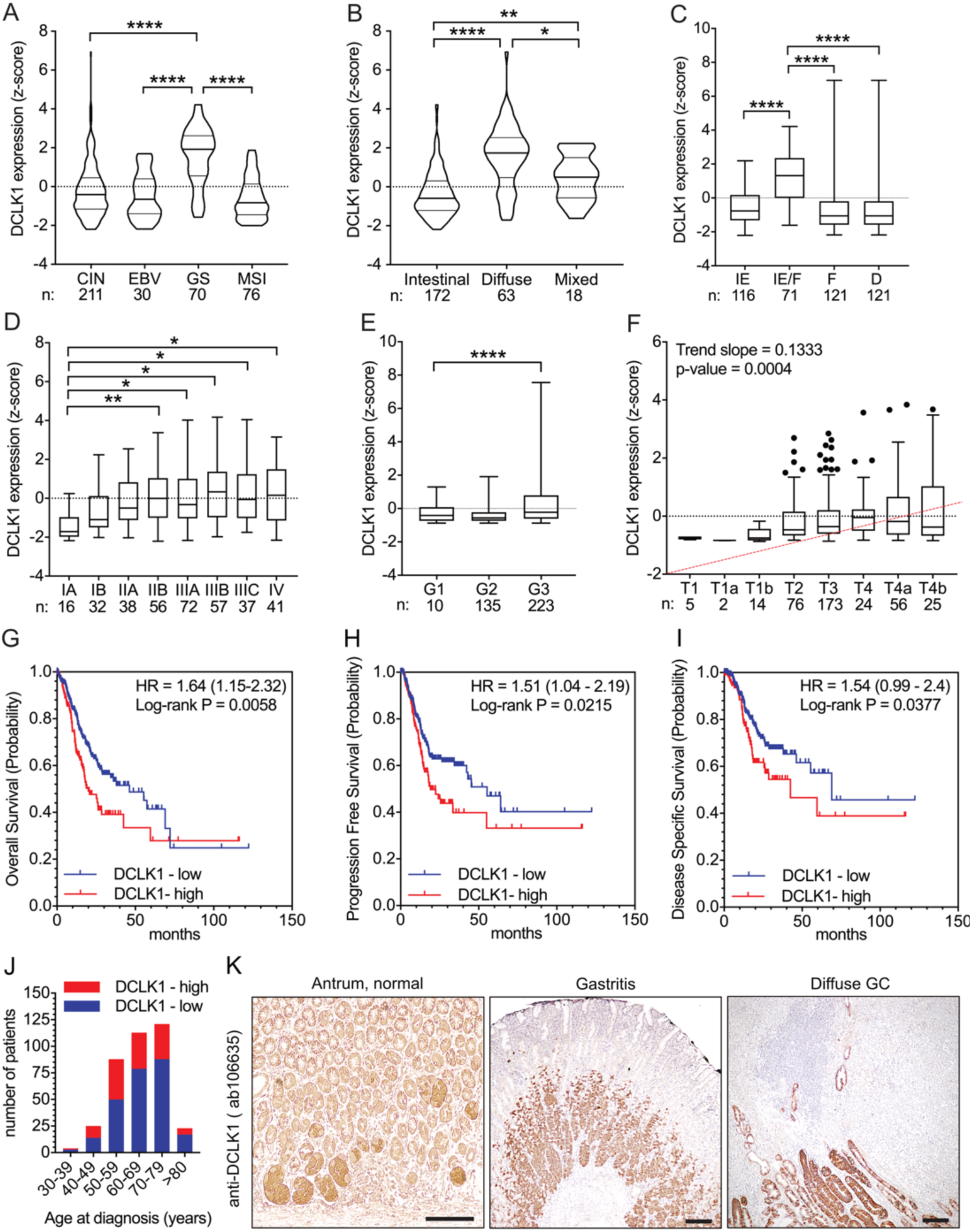
DCLK1 gene expression in gastric cancer. DCLK1 transcript levels in the TCGA-STAD dataset grouped according to (A) molecular subtype, (B) Lauren classification, (C) TME classification, (D) TNM staging, (E) neoplasm histology grade and (F) invasiveness of primary tumor classification (AJCC). (G) DCLK1-high/low Kaplan-Meier analysis for overall survival, progression-free survival (H) and disease-specific survival (I). (J) Distribution of the proportion of DCLK1-high/low tumors at patient age of diagnosis. (K) anti-DCLK1 IHC on tissue containing healthy antral stomach, gastritis or diffuse-type GC. *P < .05, **P <.01, ****P <.0001. Abbreviations: CIN, chromosomal instable; EBV, Epstein-Barr virus; GS, genomic stable; MSI, microsatellite instable; IE, Immune-Enriched; IE/F, Immune-Enriched/Fibrotic; F, Fibrotic; D, Desert; G, grade. Scale bar = 50 µm.

Next, we analyzed DCLK1 expression in 37 human gastric cancer cell lines using bulk RNAseq data from the DepMap portal (*25*). This analysis revealed low to medium mRNA expression levels of DCLK1 in the majority of GC cell lines (**Supplementary Figure 1 A**) which was comparable to the expression levels of other known cancer stem cell markers and neuronal markers but significantly lower than a selection of standard housekeeping cancer genes (**Supplementary Figure 1 B-F)**. Anti-DCLK1 immunoblot analysis showed a relatively poor correlation between DCLK1 mRNA and protein levels in the 7 GC cell lines investigated (**Supplementary Figure 1 G)**. This is in line with the moderately positive correlation (R=0.6) of RNA/protein expression for DCLK1 in cancer cell lines recently reported (*26*). Finally, we used IHC to detect DCLK1 expression in normal stomach tissue and tissues from patients with gastritis and diffuse GC. While DCLK1 positivity in the healthy stomach was confined to subsets of epithelial cells in deep antral glands, this expanded considerably in the inflamed stomach and was readily detected in the majority of cancer cells in a diffuse gastric cancer (**Figure 1 K**). Together, this data supports a pro-tumorigenic role for DCLK1 in the progression of GC.

### DCLK1 promotes growth of gastric cancer xenografts

In order to study the impact of DCLK1 expression on gastric cancer growth, we enforced expression of DCLK1 into the human gastric cancer cell line MKN1, which retains low E-cadherin, mutant TP53, a prominent EMT expression signature and other hallmarks of the GS molecular subtype (*27*). We confirmed over-expression of stably transfected full-length isoform 1 (UniProt ID: O15075-2) by immunoblotting of pools in resulting MKN1^DCLK1^ cells (**Supplementary Figure 2 A**).

We then engrafted MKN1^DCLK1^ cells or their naïve parental MKN1^naïve^ counterparts into the flanks of immuno-compromised hosts and observed that MKN1^DCLK1^ xenografts grew larger than MKN1^naïve^ xenografts despite similar tumor take-rates (18/24 vs 19/24) and proportions of proliferating and apoptotic cells (**Supplementary Figure 2 B, C, D; data not shown**). However, MKN1^DCLK1^ xenografts consistently stained stronger for vimentin and α-smooth muscle actin (α-Sma) than MKN1^naïve^ xenografts and were composed of more host derived stromal cells (**Supplementary Figure 2 E, F, G**). Likewise, differential qPCR analysis with human and mouse specific primer pairs confirmed the murine origin of the collagen-rich extracellular matrix (ECM) of MKN1^DCLK1^ tumors (**Supplementary Figure 2 H)**. We also detected significantly higher abundance of periostin, a matricellular protein expressed by mesenchymal cells, and expression of which was recently associated with a stromal signature that predicts overall worse survival in CRC patients (*28*) and of the activated isoform of the TGFβ signal transducer p-Smad2 (**Supplementary Figure 2 I**). The latter observation correlated with higher TGFβ expression in human cancer associated fibroblasts (CAFs) cultured in conditioned medium prepared from supernatant of MKN1^DCLK1^ than of MKN1^naïve^ cells (**Supplementary Figure 2 J)**. Together this data indicates that DCLK1 expression in MKN1 tumor cells promotes cancer cell extrinsic effects, resulting in host cell infiltration of tumors and coercing their production of TGFβ and associated remodeling of the tumor stroma. There is accumulating evidence in preclinical mouse models that nerves play a tumor-promoting role in GC and given that DCLK1 is a marker of neurons/neural progenitors in mice, we wanted to investigate the abundance of neurons within the TME of MKN1^naïve^ and MKN1^DCLK1^ xenografts and their overlap with DCLK1 expression. Co-immunofluorescence staining of xenograft tumors with the neuronal marker NeuroD1 and DCLK1 detected a consistently low level of less than 1% of tumor area positive for NeuroD1 with NeuroD1 positivity unaffected by DCLK1 overexpression (**Supplementary Figure 2 K,L**). Overall, we observed very little colocalization of DCLK1 and NeuroD1 which suggested that DCLK1 is not strongly expressed in cancer-associated NeuroD1+ neurons and its localization mostly restricted to MKN1 cancer cells (**Supplementary Figure 2 K,L**). Moreover, the correlation graph of DCLK1 and NeuroD1 expression in the TCGA STAD dataset indicates a weak positive correlation of R2 = 0.111, implying that low overlap of these two markers is a general feature in gastric cancers (**Supplementary Figure 2 M**).

### Selective inhibition of DCLK1 kinase activity restricts xenograft growth and stromal remodeling

The DCLK1 protein contains a functional serine/threonine kinase domain (*29*). In order to investigate whether the kinase activity of DCLK1 is required for the remodeling of the tumor stroma, we treated Balb/C hosts carrying xenograft tumors with a specific, selective and potent DCLK1 kinase inhibitor DCLK1-IN-1 (DCLK1i) (*30, 31*). Three weeks later, we observed significantly smaller tumors in the cohort of DCLK1i-treated hosts (**Figure 2 A, B**). Again this was not caused by changes in either tumor cell proliferation or apoptosis (**Supplementary Figure 3 A - D**). Instead, we observed markedly reduced staining for the stromal markers α-Sma, and Vimentin (**Figure 2 C - F**). Likewise, picrosirius red staining was diminished highlighted by the presence of fewer and thinner collagen fibers. The latter observation coincided with reduction in the collagen crosslinking enzyme lysyl oxidase, rather than of p-SMAD2 accumulation and associated TGFβ signaling, suggesting that DCLK1i-treatment impinges on TGFβ-independent tissue remodeling processes (**Figure 2 G - J**) (**Supplementary Figure 3 E, F**).

**Fig. 2.**
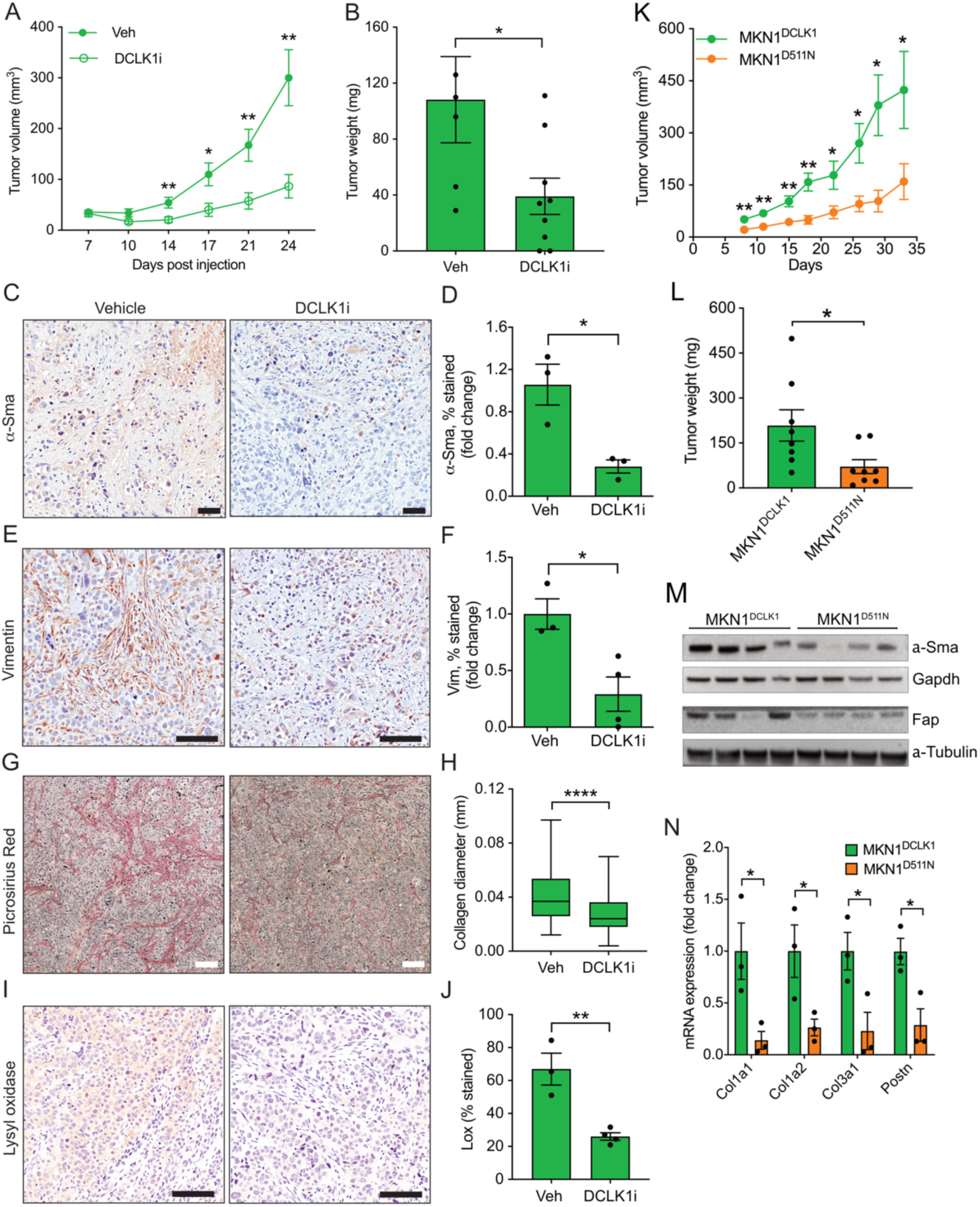
DCLK1 remodels the ECM in a kinase-dependent manner. (A) Xenograft tumor growth of MKN1^DCLK1^ cells in Balb/c nude mice treated with DCLK1 inhibitor (DCLK1i) or vehicle (Veh). (B) Xenograft tumor weights at time of harvest. (C, E, G, I) Immunohistochemical stains of xenograft tumor tissue using anti-vimentin (Vim) and α-Sma antibodies, picrosirius red stain and anti-Lysyl oxidase (Lox) antibodies. (D, F, H, J) Quantification of IHC stains. (K) Xenograft tumor growth of MKN1^DCLK1^ and MKN1^D511N^ cells in Balb/c nude mice. (L) Xenograft tumor weights at time of harvest. (M) Immunoblot of MKN1^DCLK1^ and MKN1^D511N^ xenograft tumor lysates for detection of α-Sma and Fap. Gapdh and α-tubulin are used as loading controls. (N) RT-qPCR analysis of xenograft tumors of MKN1D511N and MKN1DCLK1 cells using primers specific for the indicated murine genes. *P < .05, **P <.01, ****P <.0001. Scale bar = 100 µm.

In order to further assess the functional importance of the DCLK1 kinase domain and the on-target activity of the DCLK1 inhibitor, we engineered a DCLK1 mutant protein containing a single amino acid substitution (D to N) at the catalytic aspartate at position 511 which renders the kinase domain nonfunctional (*29*) and generated pools of stably transfected MKN1 cells (MKN1^D511N^). Similar to the effect on tumor growth and tumor burden after treatment with the DCLK1 inhibitor, MKN1^D511N^ cancer xenografts showed reduced tumor growth (**Figure 2 K, L**) and reduced expression of stromal cell markers α-Sma and Fap and ECM components collagen and periostin (**Figure 2 M, N**) (**Supplementary Figure 3 G, H**). Together this provides further support of the on-target specificity of DCLK1i and underpins the conclusion that the stromal remodeling phenotype is a direct consequence of cancer cell-extrinsic properties mediated by the kinase activity of DCLK1. In addition, we employed the kinase dead mutant to investigate whether the kinase activity also regulated cell intrinsic functions. As DCLK1 has been shown to induce cancer cell EMT and impact cancer cell stemness, we performed limiting dilution stemness assays and in vitro cell migration experiments and compared the ability of MKN1^naïve^, MKN1^DCLK1^, MKN1^D511N^ cells to grow in ultra-low attachment conditions and to migrate across a transwell membrane. While MKN1^DCLK1^ cells showed a significantly increased stem cell potential compared to the parental MKN1^naïve^ cells, this was abolished in cells expressing the kinase dead mutant D511N or in MKN1^DCLK1^ cells exposed to the DCLK1 inhibitor (**Supplementary Figure 3 I**). Similar trends to above results were observed when MKN1^naïve^ cells treated with the DCLK1 inhibitor were grown under low attachment conditions in vitro or as xenografts in vivo, yet while the trend of stemness data failed to reach significance, xenograft growth of inhibitor treated MNK1^naive^ tumors was significantly reduced, if only slightly (**Supplementary Figure 3 J, K**). These results are in keeping with the low but detectable expression of DCLK1 in naïve MKN1 cells (see **Supplementary Figure 1 G**) underlining that endogenous DCLK1 mediates the same processes as observed after its forced expression. Consistent with the literature, DCLK1 expression increased the migratory potential of MKN1^DCLK1^ cells compared to the naïve control cells (**Supplementary Figure 3 J**) and this was strongly inhibited by the DCLK1 inhibitor and almost completely abolished in MKN1^D511N^ cells (**Supplementary Figure 3 L**). Very similar results were obtained in AGS human gastric cancer cells genetically engineered to express wildtype or kinase dead mutant DCLK1, indicating that these results are not cell-type specific (**Supplementary Figure 3 M**). Together these results demonstrate that the DCLK1 kinase domain functionally regulates both cell intrinsic and extrinsic phenotypes, both of which are reversible by a recently developed specific DCLK1 kinase small kinase inhibitor.

### DCLK1 regulates ECM remodeling via kinase-dependent and independent mechanisms

Because of the known capacity of DCLK1 to promote EMT in tumor cells, we next quantified the contribution of human tumor cells to the overall pool of tumor-associated cells with stromal appearance. Accordingly, we used antibody staining for a human mitochondrial protein to discriminate between human MKN1 tumor-derived cells with fibroblast-like appearance and counterparts from the murine hosts. We found that the stroma associated with MKN1^naïve^ xenografts remained almost entirely composed of cells of human tumor origin (**Figure 3 A**). By contrast, we attributed the larger abundance of stromal cells in MKN1^DCLK1^ xenografts to an increase of both human and mouse cells (**Figure 3 A - C**). Furthermore, DCLK1i-treatment of the host reduced the proportion of human cancer cells with stromal appearance without affecting the abundance of mouse cells, suggesting that the infiltration of host cells is mediated by processes that are DCLK1 kinase-independent (**Figure 3 B, C**). The latter observations was supported by quantitative PCR analysis using primers for mouse *Hprt* housekeeping gene, which confirmed equal abundance of murine transcripts between vehicle and inhibitor treated MKN1^DCLK1^ xenografts samples (**Figure 3 D**). We then focused on human SNAI1 expression as a likely instigator of the EMT process in cancer cells and observed upregulated expression in MKN1^DCLK1^ xenografts, which was reverted in xenografts recovered from DCLK1i-treated hosts (**Figure 3 E**). Collectively, our results suggest dichotomous functions of DCLK1 in tumor cells, where DCLK1 kinase activity induces a mesenchymal transition, while the influx of host-derived stromal cells into the tumor is regulated in a kinase-independent manner.

**Fig. 3.**
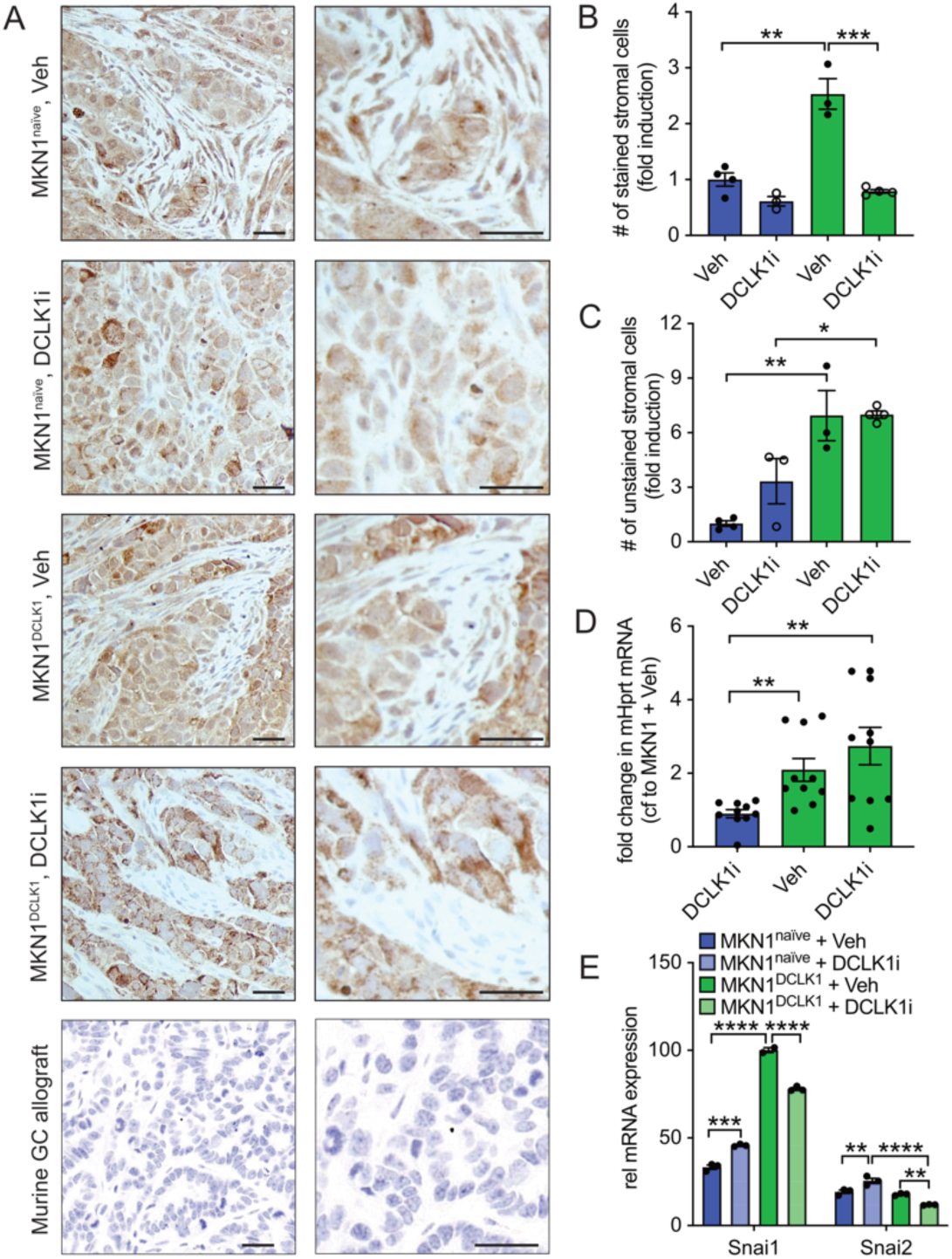
DCLK1 regulates stromal remodeling via kinase-dependent and independent mechanisms. (A) Immunohistochemical stain of xenograft tumor tissue using anti-human mitochondrial protein antibody. Bottom sample of murine GC allograft serves as antibody specificity control. (B) Quantification of stained stromal cells (fold change compared to parental vehicle control). (C) Quantification of unstained stromal cells (fold change compared to parental vehicle control). (D) Fold change in the transcript levels of murine Hprt compared to vehicle-treated MKN1^naïve^ xenograft tumors. (E) RT-qPCR analysis of EMT transcription factors Snai1 and Snai2 in xenograft tumors of MKN1^naïve^ and MKN1^DCLK1^ cells treated with vehicle or DCLK1 inhibitor. Scale bar = 50 µm. *P < .05, **P <.01, ***P < .001, ****P <.0001.

### CXCL12 chemokine promotes DCLK1-dependent EMT

The DCLK1 kinase-dependent tissue remodeling observed in the MKN1^DCLK1^ xenograft tumors implied tumor cell-extrinsic mechanisms most likely brought about by secreted mediators. We therefore used an array assay to detect chemokines in supernatants harvested from MKN1^naïve^ and MKN1^DCLK1^ cells grown in the absence or presence of the DCLK1i. We reasoned that the expression of any chemokine implicated in stromal remodeling would be increased in the supernatant of MKN1^DCLK1^ cells and decreased in DCLK1i-treated cells. Among the many mediators increased in cultures of MKN1^DCLK1^ cells that were also sensitive to DCLK1i-treatment, we detected CXCL12, Midkine and the T-cell chemoattractant CXCL16 (*32*) (**Figure 4 A**; **Supplementary Figure 4 A**). As the xenografts were grown in immunocompromised Balb/C nude mice which lack T cells, we did not consider the T cell chemoattractant CXCL16 any further (*32*). Our subsequent parsing of CXCL12 and Midkine for their capacity to mediate biological outcomes of DCLK1 based on correlating gene expression in the TCGA STAD dataset revealed a positive correlation between CXCL12 and DCLK1 (R^2^ = 0.713; p <2.2e-16), and a negative correlation between Midkine and DCLK1 (R^2^ = -0.274; p<5.611e-08) (**Figure 4 B; Supplementary Figure 4 B**). We therefore focused on the role of CXCL12 for the remaining study.

**Fig. 4.**
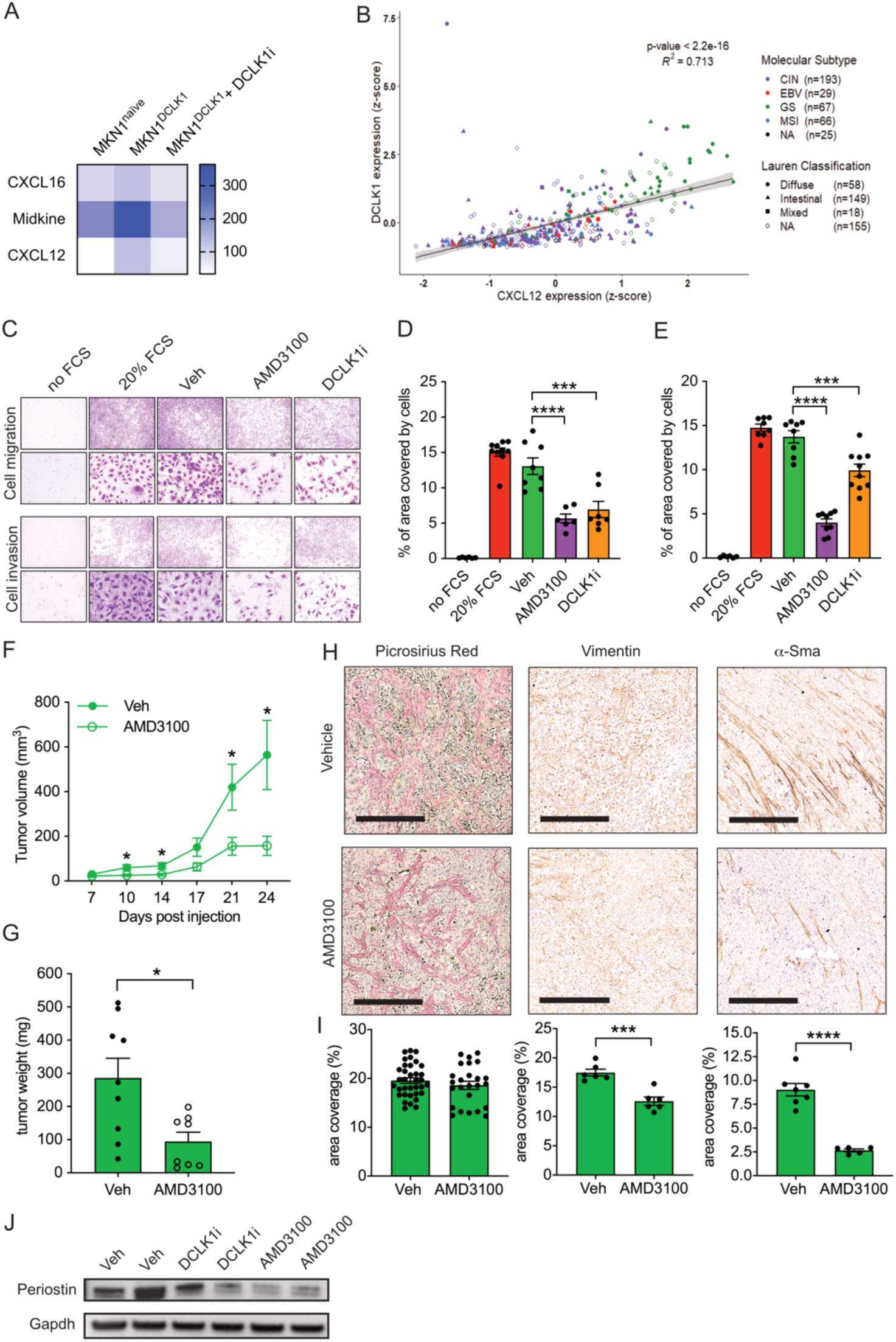
DCLK1 promotes EMT via CXCL12 chemokine. (A) Heatmap showing expression levels of selected chemokines in cell supernatants collected from cultured MKN1^naïve^ and MKN1^DCLK1^ cells and MKN1^DCLK1^ cells grown in the presence of DCLK1 inhibitor for 72 hours. (B) Correlation analysis of DCLK1 and CXCL12 expression (z-score, log2 transformed) in gastric cancers from the TCGA STAD dataset. Each dot represents a single cancer indicating both its molecular subtype and Lauren classification. (C) Cell migration and cell invasion assays using transwell inserts with MKN1^naïve^ and MKN1^DCLK1^ cells in the presence of CXCR4 antagonist (AMD3100), DCLK1 inhibitor (DCLK1i) or vehicle (Veh). No FCS is negative control where the FCS is omitted from the lower culture chamber. 20% FCS is the positive control. Images were taken using the 4x objective lens (top row) or 40x lens (bottom row). (D) Quantification of cell migration assays. (E) Quantification of cell invasion assays. (F) Xenograft tumor growth of MKN1^DCLK1^ cells in Balb/c nude mice treated with AMD3100 or vehicle (Veh). (G) Weight of xenograft tumors at time of harvest. (H) Immunohistochemical stains of xenograft tumor tissue using anti-Vimentin and α-Sma antibodies and picrosirius red stain to visualize collagen. (I) Quantification of IHC stains. (J) Immunoblot of MKN1^DCLK1^ xenograft tumor lysates for detection of Periostin and Gapdh. Scale bar = 300 µm. *P < .05, ***P < .001, ****P <.0001.

To functionally confirm a contribution of CXCL12 to DCLK1-regulated phenotypes, we first established that the abundance of its cognate CXC chemokine receptor 4 (CXCR4) remained unchanged between MKN1^DCLK1^ and MKN1^naïve^ cells. RT-qPCR, immunoblotting and anti-CXCR4 flow cytometry revealed consistent but low expression levels of the CXCR4 receptor in MKN1^naïve^ and MKN1^DCLK1^ cells (**Supplementary Figure 4 C - E**). In light of the known role of CXCL12 as a strong inducer of cell migration, we asked whether CXCL12 acted as mediator of cell motility downstream of DCLK1 by using the CXCR4-specific inhibitor AMD3100. We observed that AMD3100 inhibited *in vitro* migration and invasion of MKN1^DCLK1^ and MKN1^naïve^ cells to a level comparable to that observed with DCLK1i (**Figure 4 C, D, E**). This suggested that excessive DCLK1 activity induces mesenchymal transition in part through CXCL12. Indeed, akin to the DCLK1 inhibitor, treatment of MKN1^DCLK1^ cells with AMD3100 reduces expression of the mesenchymal marker SNAI1, while expression of the epithelial marker E-cadherin increases (**Supplementary Figure 4 F**). We then confirmed these observations in vivo by observing reduced tumor size of MKN1^DCLK1^ xenografts in hosts treated with AMD3100 (**Figure 4 F, G**). Moreover, staining with anti-α-Sma and anti-Vimentin antibodies for the detection of mesenchymal cells alongside anti-Periostin immunoblots consistently revealed a reversal of the stromal phenotype in tumors of AMD3100-treated hosts that was comparable to experiments with DCLK1i-treatment (**Figure 4 H, I, J**). However, it is likely that there is no linear signaling relationship between DCLK1 and CXCL12, because we observed that CXCL12, unlike inhibition of DCLK1, reduced tumor cell proliferation, but did not lead to reduction of stromal collagen (**Figure 4 H, I; Supplementary Figure 4 G, H**). We reasoned that these differences were likely mediated by inhibition of stromal CXCL12/CXCR4 signaling as was recently reported (*33*). In order to better ascertain this, we investigated the distribution of CXCR4 among the stromal cell populations of the MKN1^DCLK1^ tumor xenografts by flow cytometry. The results indicated wide-spread expression of CXCR4 among endothelial cells and cancer associated fibroblasts (CAFs) and to a lesser extent in macrophages and myeloid derived suppressor cells (MDSCs) supporting our evidence of DCLK1-independent effects of AMD3100 (**Supplementary Figure 4 I**).

### DCLK1 accelerates invasion and metastasis in an orthotopic model of GC

Based on the capacity of excessive DCLK1 activity to promote mesenchymal transition and stromal remodeling in vivo, we next investigated the ability of DCLK1 to promote invasion and metastasis in an orthotopic gastric cancer xenograft model (*34*). For this, we injected luciferase-tagged MKN1^DCLK1^ and MKN1^naïve^ cells into the gastric subserosa and assessed metastatic outgrowth by weekly bioluminescent imaging over 6 weeks. We observed a moderate “take rate” of MKN1^naïve^ cells (2/6 mice), which never developed metastases. Meanwhile the MKN1^DCLK1^ cells formed tumors in every host, and in 5/6 mice also developed lung metastases (**Figure 5 A, B**). Importantly, the primary MKN1^DCLK1^ cell-derived gastric tumors were also characterized by increased abundance of collagen in the ECM and increased Vimentin expression corroborating our observations with the subcutaneous xenografts (**Figure 5 C, D**). Consistent with the increased metastatic potential of MKN1^DCLK1^ cells, we also observed enhanced invasive behavior as revealed by staining with a human mitochondrial specific antibody. 83% (5/6) of primary MKN1^DCLK1^-tumors had breached the muscularis mucosa and invaded into the gastric mucosa compared to only 50% (1/2) of primary orthotopic tumors that emerged from the parental MKN1^naïve^ cells (**Figure 5 E, F**). On balance, these results illustrate that DCLK1 expression has the capacity to bestow cancer cells with pro-invasive and pro-metastatic properties.

**Fig. 5.**
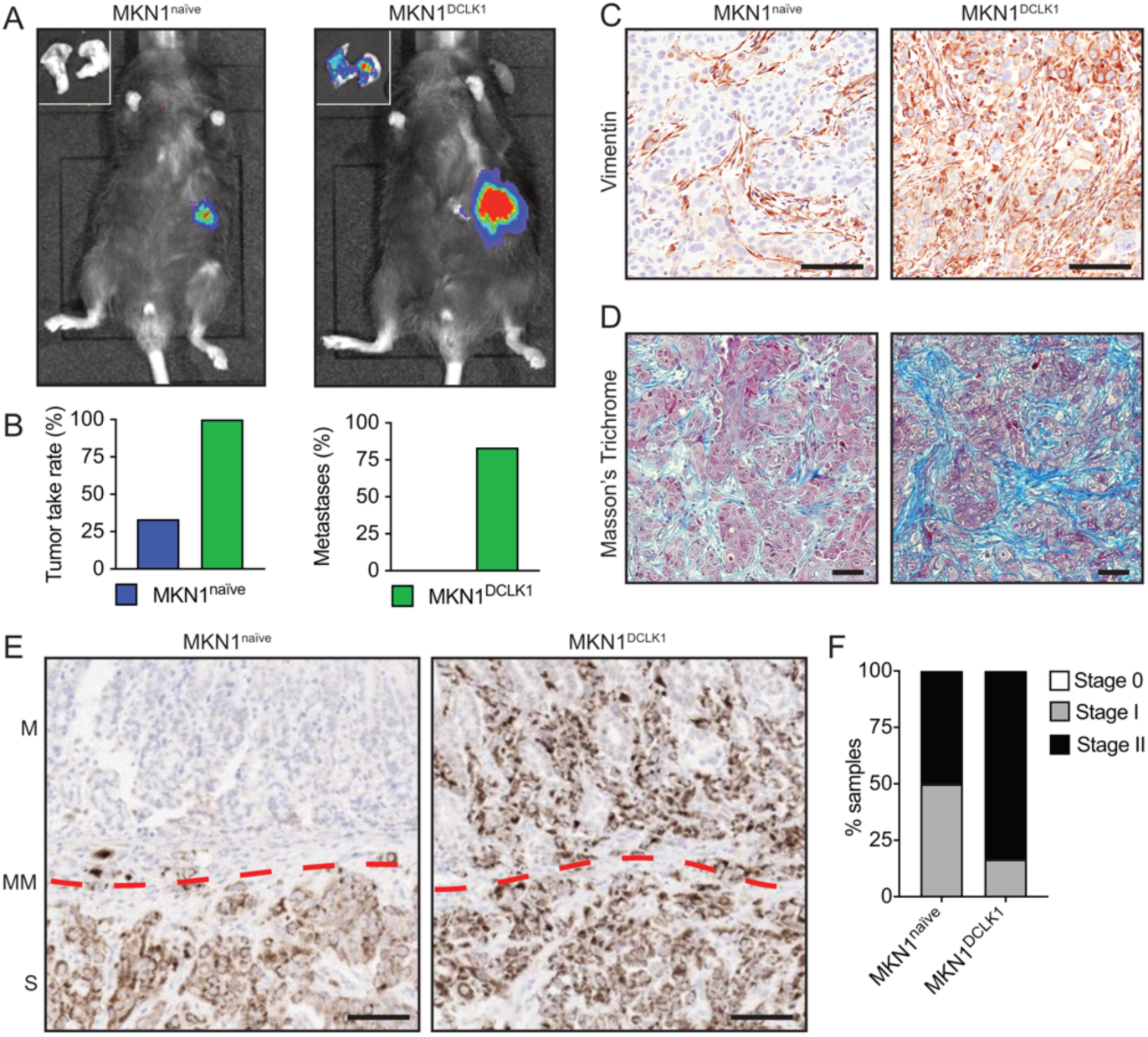
DCLK1 kinase activity accelerates invasion and metastasis in vivo. (A) Representative image of a BLI scan 6 weeks post orthotopic implantation of MKN1^naïve^ or MKN1^DCLK1^ cells. The small insets in top left corners show dissected lung lobes. (B) Tumor take-rate and the subsequent formation of lung metastases for transplanted MKN1^naïve^ and MKN1^DCLK1^ cells. IHC stains of gastric orthotopic xenografts with (C) anti-Vimentin antibody, (D) Masson’s Trichrome stain and (E) anti-human mitochondrial protein antibody for scoring of invasion stage. The dashed red line indicates the muscularis mucosa (MM). M, mucosa; S, submucosa. (F) Quantification of stage of invasion in gastric orthotopic xenografts sections stained with anti-human mitochondrial protein. Scale bar = 100 µm.

### Epithelial and stromal expression of DCLK1 in human GC

In order to extend these observations, we stained DCLK1 on a tissue microarray containing biopsies from 122 diffuse-type GC. To assist with the identification of the stromal and tumor compartments, we also stained each tumor core with H&E, anti-CD45 and anti-pan-cytokeratin antibodies (**Supplementary Figure 5 A**). Each tumor core was then scored for DCLK1 according to staining intensity (0–3) per tumor compartment (none, epithelium, stroma or both) and percentage of area stained (None, <25%, 25-50%, 50-75%, >75%) (**Figure 6 A - C, E - H**). While 18% lacked any DCLK1 immuno-reactivity, 42% of biopsies stained positive for DCLK1 in both stromal and epithelial compartments (**Figure 6 B, C**). On average, we observed more extensive areas of staining in the epithelium than in the stroma, with 30% and 17% of biopsies showing DCLK1 immuno-reactivity in at least half of all epithelial and stromal cells, respectively (**Figure 6 F, H**). These observations were reiterated in terms of staining intensity (**Figure 6 E, G**), and are reflected in the corresponding histological score (Hscore) index (**Figure 6 D**). Moreover, extending these observations to all histological GC subtypes, we performed correlation analyses between DCLK1 and the stromal fibroblast marker PGDGRA and the cancer (stem) cell marker CD44 using the TCGA STAD data. Consistent with the TMA analysis, DCLK1 positively correlated with both cancer cell and fibroblast markers, albeit with a slightly higher degree for PDGRFA (**Supplementary Figure 5 B,C)**. However, these analyses do not answer whether stromal DCLK1+ cells are a consequence of cancer cell EMT or the result of de novo expression of DCLK1 in other stromal cells. To address this question, we made use of an in house murine gastric adenocarcinoma model (*35*) whereby a transgenic *Tff1CreERT2* drives induction of oncogenic *PI3K* and *Kras* alleles as well as the expression of stabilized mutant of *p53*. These mice develop invasive gastric cancer within 3 months of induction (*MFE, MB, ME, unpublished*). Importantly, Dclk1 expression is not enforced in this GC model and cancer cells can be identified by anti-p53 stains which recognize mutant p53. Furthermore, we used Snai1 expression as a marker for cancer cell EMT. We therefore co-stained stomach sections of invasive gastric cancer with either Dclk1 and Snai1 or Dclk1 and p53 and observed broad Dclk1 positivity in cells with stromal morphology which overlapped extensively with Snai1 positive cells or p53 positive cells, respectively (**Supplementary Figure 5 D**). Finally, we conducted a surrogate experiment and stimulated cancer-associated fibroblasts in vitro with TGF-β to induce an EMT response. However, while we observed weak expression levels of DCLK1 in CAFs, TGF-β stimulation failed to induce DCLK1 expression further suggesting that DCLK1 positive stromal cells are mainly derived from cancer cells (**Figure 6 I**). Together these results strongly suggest that stromal DCLK1 expression is the result of cancer cell EMT.

**Fig. 6.**
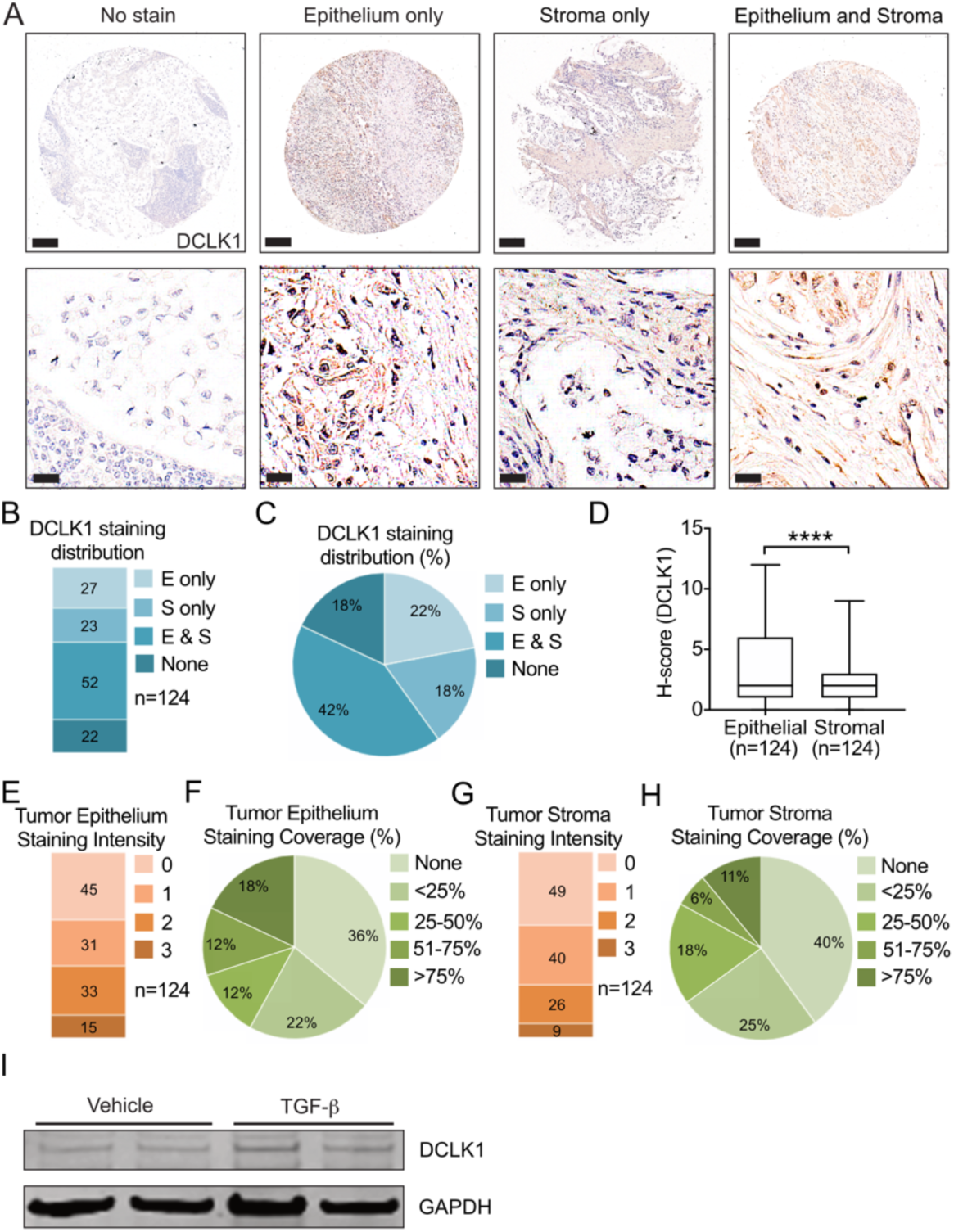
Epithelial and stromal expression of DCLK1 in human diffuse-type GC. (A) Representative photomicrographs of DCLK1 stained tissue sections of a human diffuse-type GC tissue array showing examples of gastric cancers with no DCLK1 expression (left column), DCLK1 expression restricted to epithelial cells (2nd column), DCLK1 expression restricted to stromal cells (3rd column) or DCLK1 expression observed in both stromal and epithelial cells (right column). Quantification of the staining distribution (B) and the % staining distribution (C) of DCLK1 in human diffuse-type GC tissue array. (D) Histology score (H-score) for DCLK1 in gastric cancer epithelium and stroma. H-score = Staining Intensity x % Staining Coverage. Quantification of the staining intensity (E) and the % staining coverage (F) of DCLK1 in the tumor epithelium of 124 primary diffuse-type GC tissues. (G) Quantification of the staining intensity (G) and the % staining coverage (H) of DCLK1 in the tumor stroma of 124 primary diffuse-type GC tissues. (I) Immunoblot for the detection of DCLK1 and GAPDH after stimulation of human CAFs with TGF1 (5ng, 24 hrs). Scale bars = 500 µm (top row) and 50 µm (bottom row). ****P <.0001.

This conclusion is further supported by our unsupervised clustering analysis of the TCGA STAD dataset for DCLK1 and genes associated with EMT, tumor stroma or immune signatures (*36, 37*). The heatmap indicated that DCLK1 clusters very closely with CXCL12 and with genes belonging to EMT and stromal signatures which represent the clusters most enriched in the genomic stable and diffuse subsets of GC (**Figure 7A**). As implied by our previous correlation analysis (**Supplementary Figure 4 B**), Midkine (MDK) did not co-cluster with DCLK1 while CXCL16 fell just outside an area containing genes strongly enriched in the GS and Lauren diffuse subtypes (boxed area) (**Figure 7 A**). Go-term and Reactome pathway analyses of the genes within the boxed area further emphasize processes associated with cellular migration and changes to the tumor matrix (**Supplementary Figure 6 A, B**). Given this strong association of DCLK1 with EMT and tumor stroma, we next correlated DCLK1 expression with 29 recently reported functional gene expression signatures (fges) covering known cellular and functional TME properties (*24*). The resulting Spearman correlation matrix clearly indicated highly significant positive correlations of DCLK1 with the cancer cell EMT signature as well as with angiogenic and fibrotic TME signatures across all GC subtypes (**Figure 7 B**). Moreover, we found widespread positive correlations with various fges in both the anti- and pro-tumor microenvironment categories, notably pro-tumor cytokines, macrophage/DC traffic, B and T cells, albeit, none of them were significant across all GC subtypes (**Figure 7 B**). Finally, the proliferation rate signature produced the most significant negative correlation values across all GC subtypes, corroborating our findings that DCLK1 expression did not promote proliferation of MKN1 cancer cells (**Figure 7 B, Supplementary Figure 3 A, B**).

**Fig. 7.**
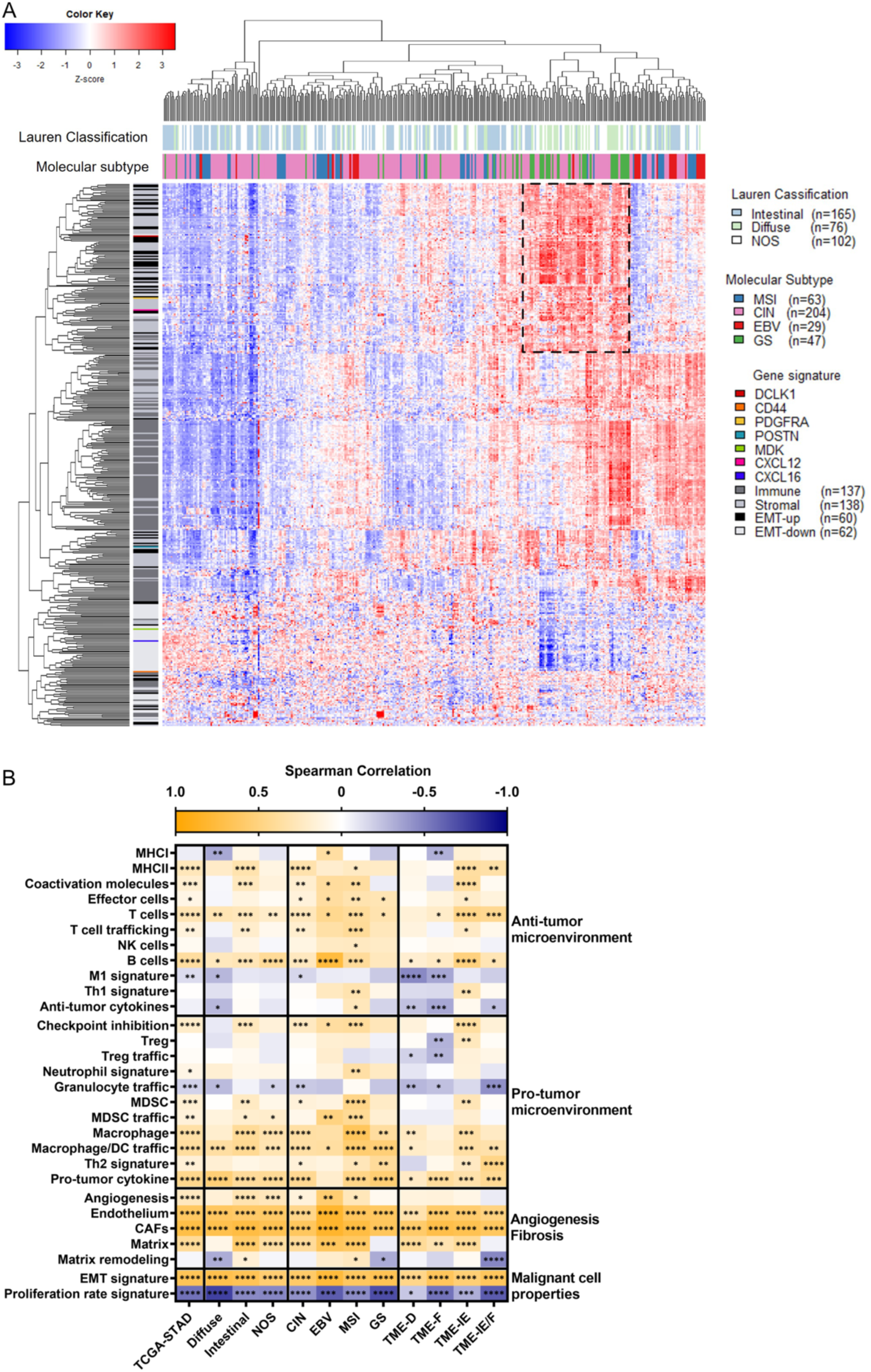
DCLK1 clusters with CXCL12, mesenchymal and stromal enriched GCs in human gastric cancer datasets. (A) Heat map of the unsupervised clustering of DCLK1 and genes associated with EMT, tumor stroma and immune response in TCGA STAD dataset separated by molecular subtype and Lauren classification (top). The position of DCLK1, CD44, PDGFRA, POSTN, CXCL12, CXCL16 and MDK genes are highlighted in various colors (left). Dashed box indicates the position of the highly enriched genes in the GS/Lauren subtype. (B) Spearman correlation between DCLK1 gene expression and 29 functional TME and malignant cell signatures subdivided by molecular, histological or TME subtypes (bottom). The individual functional gene signatures covering known cellular and TME properties are listed on the left and right y-axis, respectively. Statistical significance is indicated in each square. *P < .05, **P < .01, ***P < .001, ****P <.0001.

Collectively, our functional observations across complementing preclinical *in vitro* and *in vivo* models strongly argues for a tumor promoting role for DCLK1 kinase through the induction of a robust cancer cell EMT and the establishment of an amplified tumor secretome which simultaneously elicits paracrine and autocrine signaling pathways. As a consequence, this brings about a fibrotic TME rich in stromal infiltrates which ultimately leads to tumor progression and facilitates the formation of distant metastases.

## Discussion

During the last decade, genomic and functional analyses have supported both cancer cell intrinsic and extrinsic roles for DCLK1 as a driver of gastric tumorigenesis (*15-18, 22, 23, 38-42*). Our study presented here, unifies these different aspects of DCLK1 biology into a cooperative single mechanism.

We report here that overexpression of DCLK1 in a human gastric cancer cells drives a DCLK1 kinase-dependent transition of cancer cells to a mesenchymal phenotype and an associated surge in tumor-infiltrating host cells as the underlying mechanisms that create a fibrotic TME conducive for cancer cell invasion and metastasis. Thus, our data functionally underpins a large body of literature indicative of a positive correlation between elevated DCLK1 expression and progressive disease and reduced overall survival of patients with GC and various other solid cancer types (*43*).

Our study functionally links cancer cells with excessive DCLK1 kinase activity with their capacity to produce the chemokine CXCL12 which is widely overexpressed across many different solid tumor types (*44, 45*), and promotes tumor angiogenesis as well as the recruitment of bone marrow-derived cells to the TME. Our mechanistic insights corroborate previous observations in GI cancers, made *in silico*, of a positive correlation between expression of DCLK1 and CXCL12, TGFβ1 alongside their respective receptors and an activated TME characterized by predominance of M2 macrophages (*22*). The major source of CXCL12 in solid cancers are CAFs which exhibit properties of myofibroblasts (*33, 46, 47*). In our xenograft tumor model, CXCL12 is expressed by cancer cells and inhibition of CXCL12 action blocks the expansion of vimentin and α-Sma-positive cells in the tumor stroma, indicating that CXCL12 at least partially mediates the mesenchymal transition of cancer cells and the infiltration of bone marrow derived monocytes and/or tissue resident fibroblasts. Contrary to the DCLK1 inhibitor treatment, AMD3100-mediated inhibition of the CXCL12 receptor restrained proliferation of the MKN1^DCLK1^ tumor cells, but did not reduce the deposition of collagen. These results are reminiscent of observations in CXCL12 transgenic mice where the CXCL12/CXCR4 signaling axis promoted expansion of α-Sma-positive myofibroblasts and the proliferation of CXCR4-positive epithelial progenitor cells during gastritis, yet had minimal effect on ECM deposition and tissue fibrosis (*48*). This implies that collagen deposition is regulated by DCLK1 kinase-dependent mechanism(s) that are independent of, or bypass CXCL12 altogether. We therefore predict that therapeutic targeting of DCLK1 in these GC subtypes could translate to more profound benefits than inhibition of the CXCL12/CXCR4 signaling axis alone.

A lot of attention has been given to the role of DCLK1 as a marker of (cancer) stem cells. Notwithstanding this, our data presented here and previous studies have shown that DCLK1 expression is detectable in both tumor and stromal cells (*14, 49, 50*). Regardless of the cellular origin, mounting evidence indicates substantial trans-differentiation potential of CSCs into various stromal subtypes such as endothelial cells, pericytes, CAFs, MSCs and neurons (*51*). While such studies were mostly conducted *in vitro*, vasculogenic mimicry, the trans-differentiation of CSCs into endothelial cells, has been demonstrated *in vivo* over a decade ago (*52*). Nevertheless, cellular fusion of cancer cells with other cell types of the TME is a potential alternative pathway which could give rise to cancer cells expressing various mesenchymal, endothelial, immune or neuronal markers in vivo (*53*). Consequently, while the interrogation of publicly available single cell transcriptomic cancer datasets may identify additional DCLK1 expressing cell types and subpopulations within the TME, their exact cellular ontogeny may still remain elusive. Irrespective of the exact cellular origin of DCLK1 expressing stromal cells, considering the established role of DCLK1 in promoting cancer cell stemness and EMT of cancer cells, we conclude that at least a fraction of the stromal DCLK1+ cells are generated via EMT. This is supported by our own results which indicate that the overwhelming majority of DCLK1+ stromal cells in the tumor xenografts are of (human) cancer cell origin.

The major limiting factor of our study is the use of immunocompromised murine tumor models which precludes analysis of the impacts of DCLK1 expression and the effects of inhibiting its kinase activity on the full cellular composition of the TME. This is relevant in light of the strongly positive and negative correlations of DCLK1 gene expression with various functional signatures associated with both pro-and anti-tumourigenic properties of the TME. Further studies are therefore necessary to study DCLK1 function in immunocompetent cancer models and to gauge the impact of anti-DCLK1 therapies. The current study investigates the function of a single full-length isoform of DCLK1, however, three more splice variants have been described, and while all four isoforms contain the kinase domain, two of them are short, “kinase-only”, isoforms and lack the N-terminal DCX domains which mediate DCLK1’s microtubule-associated function (*29, 31*). Moreover, we and others have shown that the DCLK1 kinase domain negatively regulates microtubule polymerization, at least *in vitro*, and the C-terminal autoinhibitory domain (AID) of DCLK1 competes with ATP for access to the catalytic kinase domain of DCLK1, thus negatively regulating the kinase activity of DCLK1 (*29, 31, 54*). Notably, cancer-associated mutations in the AID domain lead to elevated kinase activity and reduced microtubule binding (*54, 55*). Both full-length and short isoforms have documented pro-EMT and pro-cancer functions and their kinase activity is blocked by DCLK1-IN-1 (*30, 56–58*). Nevertheless it is unclear whether kinase inhibition of short and long DCLK1 isoforms mediate anti-cancer outcomes through identical mechanisms. Future investigations such as by Liu et al. (*56*), are needed to document short and long isoform-specific cellular changes to the phosphoproteome and to identify isoform-specific kinase substrates in order to better understand the biological processes facilitated by the different DCLK1 isoforms.

The genomic stable subtype has the worst prognosis among all the gastric cancer subtypes largely due to the ineffectiveness of current therapies. Our results demonstrate that inhibition of the DCLK1 kinase activity has anti-tumorigenic potential in preclinical mouse models of GS-like stomach cancer. Intriguingly, it does so without affecting cancer cell proliferation per se, raising the possibility of combining DCLK1 small molecule inhibitors with strong anti-proliferative drug treatments to maximize anti-tumor control. Considering that the GS and diffuse GC subtypes showed the highest DCLK1 expression levels, we were intrigued to see the strong positive association of DCLK1 with EMT, fibrosis and angiogenesis across all GC cancer subtypes. This suggests that targeting DCLK1 kinase activity may not only be a novel therapeutic target for the treatment of GS and diffuse subtypes of gastric cancer but may be a novel promising pan-cancer target for stomach cancer.

## Materials and Methods

### Study Approval

All animal studies were conducted in accordance with all relevant ethical regulations for animal testing and research including the Australian code for the care and use of animals for scientific purposes. All animal studies were approved by the Animal Ethics Committee of the Ludwig Institute for Cancer Research, the Walter and Eliza Hall Institute of Medical Research, Austin Health or by the Institutional Animal Care and Use Committee at the Peter MacCallum Cancer Centre. We have complied with all relevant ethical regulations for work with human participants. Collection and usage of human gastric cancer tissues was approved by the Austin Health ethics committee and informed consent was obtained from all subjects.

### Animal Models

Bl6/Rag2/GammaC double knockout mice harboring recombinase activating gene-2 (RAG2) and cytokine receptor γ-chain (γC) mutations were bred and maintained under specific pathogen-free conditions in the research facility of the Peter MacCallum Cancer Centre. Balb/c nude mice were purchased from the Animal Resources Centre (Canning Vale, WA, Australia). All interventions were performed during the light cycle on both male and female mice. All animals had free access to water and food (standard chow). All strains were maintained on a 12-hour light/dark cycle at constant temperature. Co-housed, age- and gender-matched littermates were utilized for all experiments. The inducible *BAC(Tff1CreERT2);Kras^G12D^* model of murine GC has been reported previously (*35, 59*) and was used to generate the *BAC(Tff1CreERT2);Kras^G12D^; Pi3kca^H1047R^; p53^R172H^* mouse model of invasive gastric adenocarcinoma (MFE, MB, ME, unpublished).

### DNA Cloning

The cDNA encoding the DCLK1 isoform 1 (accession # NM_004734) was PCR amplified from a plasmid purchased from Origene (Origene Technologies, Rockville, US; Cat # RC217050) using forward primer 5’ agc aag ctt gcc acc atg tcc ttc ggc aga gac atg gag 3’and reverse primer 5’ acg gga tcc cta cat cct ggt tgc gtc ttc gtc 3’ and subcloned into pcDNA3 using HindIII and BamHI restriction sites. The construct was verified by Sanger sequencing and DCLK1 protein expression validated by immunoblotting.

### Tissue Culture

The gastric carcinoma cell lines were cultured in RPMI1640 + GlutaMax supplemented with 10% fetal calf serum (FCS) and 1% L-glutamine. They were kept in the incubator with 5% CO2 at 37°C. The cells were passaged twice a week at a 1:10 ratio.

### Western Blot

For Western Blot analysis protein lysates were prepared from tissue samples and cells. Cells were lysed in lysis buffer (10 mM Tris/HCl, 100 mM NaCl, 1% Triton X-100, 10% glycerol, 0.5 mM PMSF, 2 mM EDTA) for 30 minutes on ice after which the supernatant was collected and prepared for BCA protein concentration measurements (Pierce). 20-40µg of protein were diluted in lysis buffer, 4x loading buffer. The samples were denatured at 95°C for 5 minutes before proteins were separated according to their size on a 4-12% SDS-PAGE gels. Semi-dry blotting was used to blot the proteins on the SDS-gel onto a PVDF-membrane. The blotting step was performed with the iBlot device (Invitrogen) for 7 minutes. The membrane was blocked with blocking buffer (TBS, Odyssey Blocking Buffer, Li-COR) for 1 hour or 5% milk powder in PBS-T. The primary antibodies were diluted in blocking buffer and PBS-T (see table 2) and added to the membrane. The membrane was incubated overnight at 4°C. The next day, the membrane was washed and the secondary antibody was added in blocking buffer and incubated for 1 hour at room temperature. The membrane was washed three times for 10 minutes and subsequently dried or ECL solution (Thermo Fisher) was applied for one minute. Detection was either by fluorescence (Odyssey) or by luminescence reader (HRP, Bio-Rad).

### Real-time PCR

Total RNA was extracted from frozen xenograft tumor samples using Trizol® Reagent (Life technologies, Cat# 15596026) and cDNA was prepared from 2 µg total RNA using the high capacity cDNA reverse transcription kit (Applied Biosystems, Cat# 4368813) according to the manufacturer’s protocol. Quantitative RT-qPCR analyses were performed in technical triplicates with SensiMix SYBR kit (Bioline, Cat# QT60520) using the ViiA™ 7 Real Time PCR System (Life Technologies). Further details and the sequences of the used oligonucleotides are described in **Supplemental Table S1**.

### Chemokine Array

MKN1^naïve^ and MKN1^DCLK1^ over-expressing cells were cultured and treated with DCLK1 inhibitor (DCLK1i) and the supernatant was collected to detect the secreted chemokines. Further, protein lysates (100 µg) of MKN1^naïve^ and MKN1^DCLK1^ over-expressing cells extracted from a xenograft mouse model were analyzed for chemokine detection. For all samples, the human chemokine array kit (R&D Systems) was used according to the manufacturer’s protocol.

### Cell migration and invasion

Cell migration and invasion assays were done according to supplier’s protocol (BD Biosciences). In short, migration and invasion assay were done in individual 24 well tissue culture inserts (8µm pore size, cat # 353097). The upper chamber was plated with 4 x 10^4^ cells in serum-free medium (+/- inhibitors) while the bottom chamber contained medium with 20% FCS. 72 hours after seeding, the cells on upper chamber were wiped off and the cell on the bottom of the chamber were fixed and stained with the Diff-Quik staining solution by sequentially transferring the inserts through the fixing and two Diff-Quik staining solutions and several water rinses. The cell nuclei stain purple and the cytoplasm stains pink. The invasion assay was performed basically as described above with the exception that the top chamber was coated with Matrigel (BD Biosciences, Cat# 354578) at a final concentration of 300 µg /mL in coating buffer and incubated at 37°C for 3 hours prior to plating cells in each chamber. Stained bottom inserts were imaged with 4 x and 40 x objectives and the 40 x images were used for assessment of migration and invasion potential (% area covered by cells) using Fiji software.

### Stimulation of cancer-associated fibroblasts

Human cancer-associated fibroblasts (CAFs) were sourced as previously described (*60*) and were maintained at 37°C in 10% CO2 with in Dulbecco’s modified Eagle’s medium (DMEM)/F-12 culture media supplemented with 10% (v/v) of fetal calf serum (FCS). Cells were plated into 6 well plates at a density of 150,000 per well and stimulated for 48 hrs with 5 ng/mL human TGFβ.

Alternatively, CAFs were plated into 6-well plates at a density of 150,000 per well. MKN1naïve and MKN1DCLK1 cells were grown in 10 cm tissue culture plates until they reached 80% confluency before supernatant was collected. Dead cells were removed by centrifugation before supernatant was added to CAFs for 48 hours at which point total RNA was collected and cDNA prepared for RT-qPCR analysis using primers specific for human TGFβ.

### Immunostaining and Quantification

The paraffin-embedded tissues were dewaxed in Xylene twice for 10 minutes and subsequently rehydrated in graded ethanol baths (70%, 100%). Then the antigens were retrieved by cooking the slides in 10mM sodium citrate buffer (pH 6.0) or EDTA buffer for 18 minutes. After cooling, the endogenous peroxidases were blocked in 3% H2O2 in PBS for 20 minutes at room temperature. Nonspecific binding sites were blocked by incubating the slides in blocking buffer (5% normal goat serum (NGS) in PBS) for 60 minutes at room temperature (RT). After blocking, the sections were incubated with the primary antibody diluted in blocking buffer overnight at 4°C. The next day the sections were washed twice with 1x PBS/0.1% Tween and once with 1x PBS. Then the sections were incubated with the secondary antibody for 30 minutes at RT. After washing, the sections were incubated with ABC solution (Vectastain, ABC-Kit) diluted 1:200 in PBS for 30 minutes at RT. Sections from the human gastric cancer TMA were incubated with either anti-rabbit or mouse ImmPRESS-HRP detection kit (Vector Laboratories) for 30 minutes at RT. The sections were washed and DAB (Dako) was added onto the sections for staining. The staining reaction was stopped by applying distilled water onto the sections. The sections were then counterstained with hematoxylin and passed through ethanol gradients (100%, 70%) and xylene for dehydration. After drying, the sections were mounted with mounting media. Images were collected and analyzed with Aperio ImageScope v11.2.0.780 software (Leica Biosystems). Quantification of positive staining per µm2 was performed using an automated cell counter script in Fiji (*61*) or Aperio ImageScope nuclei detection script. Further details on the antibodies used are described in the **Supplemental Table S2**.

For immunofluorescence staining, the same rehydration and antigen retrieval protocol was used. The slides were blocked in 1% BSA in PBS with 0.5% Triton X-100 for 1 hour at room temperature. After blocking, the primary antibodies were applied and incubated overnight at 4°C. The next day the slides were washed in PBS and the secondary antibodies were applied to the slides for 1 hour at room temperature. After washing the slides in PBS, the slides were mounted with Prolong Gold anti-fade mounting solution with DAPI (Thermo Scientific). Images were acquired using a fluorescence microscope (Vectra system, Perkin Elmer) and analyzed with inForm tissue analysis software (Akoya Biosciences).

### Flow cytometry

Xenograft tumors were excised and submerged in room temperature PBS. Cells and tumor tissues were centrifuged at 100 x g at room temperature for 5 min to pellet cells and tissue pieces before being resuspended in 5ml tissue culture medium (1x HBSS, Ca+ Mg+ free).

Tissue was transferred into a 60 mm petri dish and minced with scalpel blade to obtain ∼ 1-3 mm3 pieces. Minced tissue was transferred into 15 mL conical tube and centrifuged at 100 x g for 5 min at room temperature. Pelleted tissue was resuspended in 4.7 mL warm tissue culture medium and 300 µL enzymatic digestion mix was added containing Collagenase III, Dispase and DNase I at 1 mg/mL, 0.4 U/mL and 2 µg/mL final concentrations, respectively and incubated on a nutating platform at 37°C for 60 min. The cell suspension was then triturated with a 10 ml plastic serological pipet until a homogenous cell suspension was achieved before being strained through a 70 µm cell strainer. Single cells were collected by centrifugation at 100 x g for 10 min at room temperature and supernatant was discarded. Cell pellet was resuspended in warm cell culture medium and viable cells were counted using Trypan Blue. Cells were then pellet again at 100 x g for 10 min at room temperature and resuspended in 1ml staining buffer (2.5% FCS in PBS). Cells were blocked in FcR blocking reagent (Milteny Biotec) on ice for 20 min and incubated with cell type-specific antibody panels on ice for 1 hr for the staining of endothelial cells (CD45- CD31+), CAFs (Cd45- Epcam- CD31- PDPN+ PDGFRA+), macrophages (CD45+ CD11B+) mMDSCs (CD45+ CD11B+ F480+ LY6C- LY6G-) and gMDSCs (CD45+ CD11B+ F480- LY6G+ LY6C-). SYTOX Blue (Thermo Fisher Scientific) was used for dead cell exclusion. For dilution of primary and secondary FACS antibodies see **Supplemental Table S3**. Stained cells were acquired for flow cytometry on a FACSCanto II (BD Biosciences) and analyzed using FlowJo software (BD Biosciences).

### Limiting dilution assay

MKN1^naïve^, MKN1^DCLK1^, MKN1^D511N^ cancer cells were grown to 80% confluency and then harvested by trypsinization and counted using a hemocytometer. Cells were then diluted in growth media and manually seeded into rows of 12 wells of a 96-well clear round bottom ultra-low attachment spheroid microplate at 500, 50, 10, 5 and 1 cell(s) / well and incubated at 37°C, 5% CO2 for 14 days. 24 hrs after initial seeding, the number of cells in each well was manually scored under the microscope and attributed to the following 5 categories: 500 cells/well, 10-50 cells/well, 5-9 cells/well, 2-4 cells/well and 1 cell/well. Wells without cells were excluded from further analysis. DCLK1-IN-1 was used at final concentration of 1µM. Two weeks after initial seeding, each well was scored for the presence or absence of a cancer cell spheroid.

### Histological TMA Scoring

A pathologist was consulted for the diagnosis of the samples, and scoring of immune-stained slides was performed independently by two investigators (AC, MB). The scoring was carried out based on two different parameters: (1) staining intensity and (2) amount of tissue involved. Epithelia and stroma were scored separately. The intensity was measured and scored from 0 to 3, no staining = 0, weak staining = 1, moderate staining = 2 and strong staining = 3. The amount of tissue involved was measured and scored from 0 to 4, no tissue involved (None) = 0, <25 % involved = 1, 25–50 % involved = 2, 51–75 % involved = 3 and >75 % involved = 4. Finally, the intensity score was multiplied by tissue involvement (%) score to obtain DCLK1 histology score.

### Orthotopic gastric cancer model

8-10 week old female Balb/c nude mice were subcutaneously engrafted with 5×10^6^ MKN1^naïve^ or MKN1^DCLK1^ cells in both flanks. Once tumor xenografts became visible, mice were randomly divided into the various treatment cohorts. Treatments were daily during the week with treatment holidays during the weekend. Tumor xenograft diameters were recorded three times per week by caliper. Xenograft volume was calculated according to the following formula: V = (L^2^ x W)/2. Mice were sacrificed before the tumors reached 1500 mm^3^. For orthotopic grafting, 8-10 week old Bl6/Rag2/γC double knockout mice were injected orthotopically with 5×10^4^ luciferase-tagged MKN1^naïve^ or MKN1^DCLK1^ into the gastric subserosa as previously described (*34*).

### Drug Treatments (DCLK1i, AMD3100)

MKN1^naïve^, MKN1^DCLK1^ cells were treated with 1 µM DCLK1-IN-1 (DCLK1i), AMD3100 or vehicle (0.1% DMSO) for 24 to 72 hours. For in vivo application, the DCLK1 inhibitor (DCLK1i; DCLK1-IN-1 kindly provided by (*30*)) was prepared at 3 mg/ml in 5% NMP / 5% Solutol HS-15 / 90% saline. Skin xenograft bearing Balb/c nude mice were treated daily by oral gavage at a dose of 30 mg/kg DCLK1 inhibitor or vehicle control for 3 weeks. The DCLK1 inhibitor and vehicle were prepared fresh every week and stored at 4°C. Likewise, AMD3100 was diluted in PBS at 350 µg/ml and administered at 3.5 mg/kg by daily intraperitoneal injection for 3 weeks.

### TCGA-STAD data analysis

Preprocessed TCGA-STAD RNAseq Log2(RPKM+1) values were downloaded via the Xena platform from the University of California, Santa Cruz (UCSC) (*62*), and Z-score normalized per gene row (z = (x-µ)/σ). The clinical data, including: tumor staging, Lauren classification and molecular subtype, were obtained from cbioportal (*63, 64*) and linked to the TCGA-patient-ID number. The 130 EMT-up and down regulated gene signatures, and the 280 stromal- and immune gene signatures were acquired from published pan-cancer meta-analysis (*36, 37*). DCLK1, CXCL12, CXCL16, Midkine, periostin and the EMT, stromal, and immune genes were extracted from the Z-score normalized RNAseq data set. The extracted data-frame was used for spearman-correlations, unsupervised clustering, and heatmap visualization performed in Rstudio version 4.0.0 (64bit) (*65*).

### Statistical Analysis

Statistical analyses were performed in Prism (GraphPad, San Diego, CA). Statistical significance was calculated by either 2-tailed unpaired t tests assuming equal variance or 1-way analysis of variance. Data are expressed as mean ± standard error of the mean.

## Acknowledgments

This project was supported by the National Health and Medical Research Council of Australia (NHMRC) Senior Research Fellowship (1079257), Program Grant (1092788) and Investigator Grant (1173814) to ME, and a NHMRC Project Grant (1143020) and a La Trobe RFA Understanding Disease grant to MB, and the Operational Infrastructure Support Program, Victorian Government, Australia and a DF/HCC GI SPORE Developmental Research Project Award P50CA127003 (N.S.G.) and Hale Center for Pancreatic Research (N.S.G.) We acknowledge The Collie Foundation for providing funds to purchase the Leica Aperio slide scanner. We acknowledge the Australian Cancer Research Foundation and The Collie Foundation for providing funds to purchase the Zeiss 980 confocal microscope. We acknowledge The Ian Potter Foundation for providing funds to purchase the Vectra system. We are indebted to Dr. David Williams (Department of Pathology, Austin Hospital) for providing the gastric cancer tissue arrays. We thank the members of the Cancer and Inflammation program for helpful discussions and comments.

## Author contributions

Conceptualization: MB, ME, IL

Investigation: MB, SS, AA, ROK, JT, ALEC, ALC, SF, LE, RB, PT, DB, MFE, OP

Methodology: MFE, FF, NSG, OP

Visualization: JT, ALEC, SS, MB

Supervision: MB, ME

Writing - original draft: MB, ME

Writing – review & editing: MB, ME, SS, JT, ALEC, OP, IL, FF, RB, AB, NSG

## Data and materials availability

All data and materials in the main text or the supplementary materials used in the analyses will be made available to the research community upon reasonable request.

## References

1. J. Ferlay, I. Soerjomataram, R. Dikshit, S. Eser, C. Mathers, M. Rebelo, D. M. Parkin, D. Forman, F. Bray, Cancer incidence and mortality worldwide: sources, methods and major patterns in GLOBOCAN 2012. Int J Cancer 136, E359–386 (2015).

2. N. Cancer Genome Atlas Research, Comprehensive molecular characterization of gastric adenocarcinoma. Nature 513, 202–209 (2014).

3. P. Correa, Human gastric carcinogenesis: a multistep and multifactorial process--First American Cancer Society Award Lecture on Cancer Epidemiology and Prevention. Cancer Res 52, 6735–6740 (1992).

4. N. Kemi, M. Eskuri, A. Herva, J. Leppanen, H. Huhta, O. Helminen, J. Saarnio, T. J. Karttunen, J. H. Kauppila, Tumour-stroma ratio and prognosis in gastric adenocarcinoma. Br J Cancer 119, 435–439 (2018).

5. J. Al-Bassam, R. S. Ozer, D. Safer, S. Halpain, R. A. Milligan, MAP2 and tau bind longitudinally along the outer ridges of microtubule protofilaments. J Cell Biol 157, 1187–1196 (2002).

6. S. Bechstedt, G. J. Brouhard, Doublecortin recognizes the 13-protofilament microtubule cooperatively and tracks microtubule ends. Dev Cell 23, 181–192 (2012).

7. S. Bechstedt, K. Lu, G. J. Brouhard, Doublecortin recognizes the longitudinal curvature of the microtubule end and lattice. Curr Biol 24, 2366–2375 (2014).

8. F. M. Coquelle, T. Levy, S. Bergmann, S. G. Wolf, D. Bar-El, T. Sapir, Y. Brody, I. Orr, N. Barkai, G. Eichele, O. Reiner, Common and divergent roles for members of the mouse DCX superfamily. Cell Cycle 5, 976–983 (2006).

9. C. A. Moores, M. Perderiset, F. Francis, J. Chelly, A. Houdusse, R. A. Milligan, Mechanism of microtubule stabilization by doublecortin. Mol Cell 14, 833–839 (2004).

10. C. A. Moores, M. Perderiset, C. Kappeler, S. Kain, D. Drummond, S. J. Perkins, J. Chelly, R. Cross, A. Houdusse, F. Francis, Distinct roles of doublecortin modulating the microtubule cytoskeleton. EMBO J 25, 4448–4457 (2006).

11. O. Reiner, F. M. Coquelle, B. Peter, T. Levy, A. Kaplan, T. Sapir, I. Orr, N. Barkai, G. Eichele, S. Bergmann, The evolving doublecortin (DCX) superfamily. BMC Genomics 7, 188 (2006).

12. N. Weygant, Y. Ge, D. Qu, J. S. Kaddis, W. L. Berry, R. May, P. Chandrakesan, E. Bannerman-Menson, K. J. Vega, J. J. Tomasek, M. S. Bronze, G. An, C. W. Houchen, Survival of Patients with Gastrointestinal Cancers Can Be Predicted by a Surrogate microRNA Signature for Cancer Stem-like Cells Marked by DCLK1 Kinase. Cancer Res 76, 4090–4099 (2016).

13. K. Nishio, K. Kimura, R. Amano, B. Nakata, S. Yamazoe, G. Ohira, K. Miura, N. Kametani, H. Tanaka, K. Muguruma, K. Hirakawa, M. Ohira, Doublecortin and CaM kinase-like-1 as an independent prognostic factor in patients with resected pancreatic carcinoma. World J Gastroenterol 23, 5764–5772 (2017).

14. D. Qu, J. Johnson, P. Chandrakesan, N. Weygant, R. May, N. Aiello, A. Rhim, L. Zhao, W. Zheng, S. Lightfoot, S. Pant, J. Irvan, R. Postier, J. Hocker, J. S. Hanas, N. Ali, S. M. Sureban, G. An, M. J. Schlosser, B. Stanger, C. W. Houchen, Doublecortin-like kinase 1 is elevated serologically in pancreatic ductal adenocarcinoma and widely expressed on circulating tumor cells. PLoS One 10, e0118933 (2015).

15. K. Wang, S. T. Yuen, J. Xu, S. P. Lee, H. H. Yan, S. T. Shi, H. C. Siu, S. Deng, K. M. Chu, S. Law, K. H. Chan, A. S. Chan, W. Y. Tsui, S. L. Ho, A. K. Chan, J. L. Man, V. Foglizzo, M. K. Ng, A. S. Chan, Y. P. Ching, G. H. Cheng, T. Xie, J. Fernandez, V. S. Li, H. Clevers, P. A. Rejto, M. Mao, S. Y. Leung, Whole-genome sequencing and comprehensive molecular profiling identify new driver mutations in gastric cancer. Nat Genet 46, 573–582 (2014).

16. P. Chandrakesan, N. Weygant, R. May, D. Qu, H. R. Chinthalapally, S. M. Sureban, N. Ali, S. A. Lightfoot, S. Umar, C. W. Houchen, DCLK1 facilitates intestinal tumor growth via enhancing pluripotency and epithelial mesenchymal transition. Oncotarget 5, 9269–9280 (2014).

17. W. Liu, S. Wang, Q. Sun, Z. Yang, M. Liu, H. Tang, DCLK1 promotes epithelial-mesenchymal transition via the PI3K/Akt/NF-kappaB pathway in colorectal cancer. Int J Cancer, (2017).

18. S. M. Sureban, R. May, S. A. Lightfoot, A. B. Hoskins, M. Lerner, D. J. Brackett, R. G. Postier, R. Ramanujam, A. Mohammed, C. V. Rao, J. H. Wyche, S. Anant, C. W. Houchen, DCAMKL-1 regulates epithelial-mesenchymal transition in human pancreatic cells through a miR-200a-dependent mechanism. Cancer Res 71, 2328–2338 (2011).

19. T. Gao, M. Wang, L. Xu, T. Wen, J. Liu, G. An, DCLK1 is up-regulated and associated with metastasis and prognosis in colorectal cancer. J Cancer Res Clin Oncol 142, 2131–2140 (2016).

20. G. A. Busslinger, B. L. A. Weusten, A. Bogte, H. Begthel, L. A. A. Brosens, H. Clevers, Human gastrointestinal epithelia of the esophagus, stomach, and duodenum resolved at single-cell resolution. Cell Rep 34, 108819 (2021).

21. B. Schutz, A. L. Ruppert, O. Strobel, M. Lazarus, Y. Urade, M. W. Buchler, E. Weihe, Distribution pattern and molecular signature of cholinergic tuft cells in human gastro-intestinal and pancreatic-biliary tract. Sci Rep 9, 17466 (2019).

22. X. Wu, D. Qu, N. Weygant, J. Peng, C. W. Houchen, Cancer Stem Cell Marker DCLK1 Correlates with Tumorigenic Immune Infiltrates in the Colon and Gastric Adenocarcinoma Microenvironments. Cancers (Basel) 12, (2020).

23. A. L. E. Carli, S. Afshar-Sterle, A. Rai, H. Fang, R. O’Keefe, J. Tse, F. M. Ferguson, N. S. Gray, M. Ernst, D. W. Greening, M. Buchert, Cancer stem cell marker DCLK1 reprograms small extracellular vesicles toward migratory phenotype in gastric cancer cells. Proteomics 21, e2000098 (2021).

24. A. Bagaev, N. Kotlov, K. Nomie, V. Svekolkin, A. Gafurov, O. Isaeva, N. Osokin, I. Kozlov, F. Frenkel, O. Gancharova, N. Almog, M. Tsiper, R. Ataullakhanov, N. Fowler, Conserved pan-cancer microenvironment subtypes predict response to immunotherapy. Cancer Cell 39, 845–865 e847 (2021).

25. A. Tsherniak, F. Vazquez, P. G. Montgomery, B. A. Weir, G. Kryukov, G. S. Cowley, S. Gill, W. F. Harrington, S. Pantel, J. M. Krill-Burger, R. M. Meyers, L. Ali, A. Goodale, Y. Lee, G. Jiang, J. Hsiao, W. F. J. Gerath, S. Howell, E. Merkel, M. Ghandi, L. A. Garraway, D. E. Root, T. R. Golub, J. S. Boehm, W. C. Hahn, Defining a Cancer Dependency Map. Cell 170, 564–576 e516 (2017).

26. D. P. Nusinow, J. Szpyt, M. Ghandi, C. M. Rose, E. R. McDonald, 3rd, M. Kalocsay, J. Jane-Valbuena, E. Gelfand, D. K. Schweppe, M. Jedrychowski, J. Golji, D. A. Porter, T. Rejtar, Y. K. Wang, G. V. Kryukov, F. Stegmeier, B. K. Erickson, L. A. Garraway, W. R. Sellers, S. P. Gygi, Quantitative Proteomics of the Cancer Cell Line Encyclopedia. Cell 180, 387–402 e316 (2020).

27. R. Cristescu, J. Lee, M. Nebozhyn, K. M. Kim, J. C. Ting, S. S. Wong, J. Liu, Y. G. Yue, J. Wang, K. Yu, X. S. Ye, I. G. Do, S. Liu, L. Gong, J. Fu, J. G. Jin, M. G. Choi, T. S. Sohn, J. H. Lee, J. M. Bae, S. T. Kim, S. H. Park, I. Sohn, S. H. Jung, P. Tan, R. Chen, J. Hardwick, W. K. Kang, M. Ayers, D. Hongyue, C. Reinhard, A. Loboda, S. Kim, A. Aggarwal, Molecular analysis of gastric cancer identifies subtypes associated with distinct clinical outcomes. Nat Med 21, 449–456 (2015).

28. A. Calon, E. Lonardo, A. Berenguer-Llergo, E. Espinet, X. Hernando-Momblona, M. Iglesias, M. Sevillano, S. Palomo-Ponce, D. V. Tauriello, D. Byrom, C. Cortina, C. Morral, C. Barcelo, S. Tosi, A. Riera, C. S. Attolini, D. Rossell, E. Sancho, E. Batlle, Stromal gene expression defines poor-prognosis subtypes in colorectal cancer. Nat Genet 47, 320–329 (2015).

29. O. Patel, W. Dai, M. Mentzel, M. D. Griffin, J. Serindoux, Y. Gay, S. Fischer, S. Sterle, A. Kropp, C. J. Burns, M. Ernst, M. Buchert, I. S. Lucet, Biochemical and Structural Insights into Doublecortin-like Kinase Domain 1. Structure 24, 1550–1561 (2016).

30. F. M. Ferguson, B. Nabet, S. Raghavan, Y. Liu, A. L. Leggett, M. Kuljanin, R. L. Kalekar, A. Yang, S. He, J. Wang, R. W. S. Ng, R. Sulahian, L. Li, E. J. Poulin, L. Huang, J. Koren, N. Dieguez-Martinez, S. Espinosa, Z. Zeng, C. R. Corona, J. D. Vasta, R. Ohi, T. Sim, N. D. Kim, W. Harshbarger, J. M. Lizcano, M. B. Robers, S. Muthaswamy, C. Y. Lin, A. T. Look, K. M. Haigis, J. D. Mancias, B. M. Wolpin, A. J. Aguirre, W. C. Hahn, K. D. Westover, N. S. Gray, Discovery of a selective inhibitor of doublecortin like kinase 1. Nat Chem Biol 16, 635–643 (2020).

31. O. Patel, M. J. Roy, A. Kropp, J. M. Hardy, W. Dai, I. S. Lucet, Structural basis for small molecule targeting of Doublecortin Like Kinase 1 with DCLK1-IN-1. Commun Biol 4, 1105 (2021).

32. S. Hojo, K. Koizumi, K. Tsuneyama, Y. Arita, Z. Cui, K. Shinohara, T. Minami, I. Hashimoto, T. Nakayama, H. Sakurai, Y. Takano, O. Yoshie, K. Tsukada, I. Saiki, High-level expression of chemokine CXCL16 by tumor cells correlates with a good prognosis and increased tumor-infiltrating lymphocytes in colorectal cancer. Cancer Res 67, 4725–4731 (2007).

33. M. Quante, S. P. Tu, H. Tomita, T. Gonda, S. S. Wang, S. Takashi, G. H. Baik, W. Shibata, B. Diprete, K. S. Betz, R. Friedman, A. Varro, B. Tycko, T. C. Wang, Bone marrow-derived myofibroblasts contribute to the mesenchymal stem cell niche and promote tumor growth. Cancer Cell 19, 257–272 (2011).

34. R. A. Busuttil, D. S. Liu, N. Di Costanzo, J. Schroder, C. Mitchell, A. Boussioutas, An orthotopic mouse model of gastric cancer invasion and metastasis. Sci Rep 8, 825 (2018).

35. S. Thiem, M. F. Eissmann, J. Elzer, A. Jonas, T. L. Putoczki, A. Poh, P. Nguyen, A. Preaudet, D. Flanagan, E. Vincan, P. Waring, M. Buchert, A. Jarnicki, M. Ernst, Stomach-Specific Activation of Oncogenic KRAS and STAT3-Dependent Inflammation Cooperatively Promote Gastric Tumorigenesis in a Preclinical Model. Cancer Res 76, 2277–2287 (2016).

36. C. J. Groger, M. Grubinger, T. Waldhor, K. Vierlinger, W. Mikulits, Meta-analysis of gene expression signatures defining the epithelial to mesenchymal transition during cancer progression. PLoS One 7, e51136 (2012).

37. K. Yoshihara, M. Shahmoradgoli, E. Martinez, R. Vegesna, H. Kim, W. Torres-Garcia, V. Trevino, H. Shen, P. W. Laird, D. A. Levine, S. L. Carter, G. Getz, K. Stemke-Hale, G. B. Mills, R. G. Verhaak, Inferring tumour purity and stromal and immune cell admixture from expression data. Nat Commun 4, 2612 (2013).

38. P. Chandrakesan, J. Panneerselvam, D. Qu, N. Weygant, R. May, M. S. Bronze, C. W. Houchen, Regulatory Roles of Dclk1 in Epithelial Mesenchymal Transition and Cancer Stem Cells. J Carcinog Mutagen 7, (2016).

39. Y. Ikezono, H. Koga, J. Akiba, M. Abe, T. Yoshida, F. Wada, T. Nakamura, H. Iwamoto, A. Masuda, T. Sakaue, H. Yano, O. Tsuruta, T. Torimura, Pancreatic Neuroendocrine Tumors and EMT Behavior Are Driven by the CSC Marker DCLK1. Mol Cancer Res 15, 744–752 (2017).

40. Z. Q. Liu, W. F. He, Y. J. Wu, S. L. Zhao, L. Wang, Y. Y. Ouyang, S. Y. Tang, LncRNA SNHG1 promotes EMT process in gastric cancer cells through regulation of the miR-15b/DCLK1/Notch1 axis. BMC Gastroenterol 20, 156 (2020).

41. S. Makino, H. Takahashi, D. Okuzaki, N. Miyoshi, N. Haraguchi, T. Hata, C. Matsuda, H. Yamamoto, T. Mizushima, M. Mori, Y. Doki, DCLK1 integrates induction of TRIB3, EMT, drug resistance and poor prognosis in colorectal cancer. Carcinogenesis 41, 394–396 (2020).

42. N. Weygant, D. Qu, R. May, R. M. Tierney, W. L. Berry, L. Zhao, S. Agarwal, P. Chandrakesan, H. R. Chinthalapally, N. T. Murphy, J. D. Li, S. M. Sureban, M. J. Schlosser, J. J. Tomasek, C. W. Houchen, DCLK1 is a broadly dysregulated target against epithelial-mesenchymal transition, focal adhesion, and stemness in clear cell renal carcinoma. Oncotarget 6, 2193–2205 (2015).

43. W. Shi, F. Li, S. Li, J. Wang, Q. Wang, X. Yan, Q. Zhang, L. Chai, M. Li, Increased DCLK1 correlates with the malignant status and poor outcome in malignant tumors: a meta-analysis. Oncotarget 8, 100545–100557 (2017).

44. F. Balkwill, Cancer and the chemokine network. Nat Rev Cancer 4, 540–550 (2004).

45. N. Karin, The multiple faces of CXCL12 (SDF-1alpha) in the regulation of immunity during health and disease. J Leukoc Biol 88, 463–473 (2010).

46. A. Orimo, P. B. Gupta, D. C. Sgroi, F. Arenzana-Seisdedos, T. Delaunay, R. Naeem, V. J. Carey, A. L. Richardson, R. A. Weinberg, Stromal fibroblasts present in invasive human breast carcinomas promote tumor growth and angiogenesis through elevated SDF-1/CXCL12 secretion. Cell 121, 335–348 (2005).

47. S. Tu, G. Bhagat, G. Cui, S. Takaishi, E. A. Kurt-Jones, B. Rickman, K. S. Betz, M. Penz-Oesterreicher, O. Bjorkdahl, J. G. Fox, T. C. Wang, Overexpression of interleukin-1beta induces gastric inflammation and cancer and mobilizes myeloid-derived suppressor cells in mice. Cancer Cell 14, 408–419 (2008).

48. W. Shibata, H. Ariyama, C. B. Westphalen, D. L. Worthley, S. Muthupalani, S. Asfaha, Z. Dubeykovskaya, M. Quante, J. G. Fox, T. C. Wang, Stromal cell-derived factor-1 overexpression induces gastric dysplasia through expansion of stromal myofibroblasts and epithelial progenitors. Gut 62, 192–200 (2013).

49. J. Li, Y. Wang, J. Ge, W. Li, L. Yin, Z. Zhao, S. Liu, H. Qin, J. Yang, L. Wang, B. Ni, Y. Liu, H. Wang, Doublecortin-Like Kinase 1 (DCLK1) Regulates B Cell-Specific Moloney Murine Leukemia Virus Insertion Site 1 (Bmi-1) and is Associated with Metastasis and Prognosis in Pancreatic Cancer. Cell Physiol Biochem 51, 262–277 (2018).

50. J. Whorton, S. M. Sureban, R. May, D. Qu, S. A. Lightfoot, M. Madhoun, M. Johnson, W. M. Tierney, J. T. Maple, K. J. Vega, C. W. Houchen, DCLK1 is detectable in plasma of patients with Barrett’s esophagus and esophageal adenocarcinoma. Dig Dis Sci 60, 509–513 (2015).

51. Z. Huang, T. Wu, A. Y. Liu, G. Ouyang, Differentiation and transdifferentiation potentials of cancer stem cells. Oncotarget 6, 39550–39563 (2015).

52. L. Ricci-Vitiani, R. Pallini, M. Biffoni, M. Todaro, G. Invernici, T. Cenci, G. Maira, E. A. Parati, G. Stassi, L. M. Larocca, R. De Maria, Tumour vascularization via endothelial differentiation of glioblastoma stem-like cells. Nature 468, 824–828 (2010).

53. J. M. Pawelek, A. K. Chakraborty, Fusion of tumour cells with bone marrow-derived cells: a unifying explanation for metastasis. Nat Rev Cancer 8, 377–386 (2008).

54. L. Cheng, Z. Yang, W. Guo, C. Wu, S. Liang, A. Tong, Z. Cao, R. F. Thorne, S. Y. Yang, Y. Yu, Q. Chen, DCLK1 autoinhibition and activation in tumorigenesis. Innovation (N Y) 3, 100191 (2022).

55. R. L. Agulto, M. M. Rogers, T. C. Tan, A. Ramkumar, A. M. Downing, H. Bodin, J. Castro, D. W. Nowakowski, K. M. Ori-McKenney, Autoregulatory control of microtubule binding in doublecortin-like kinase 1. Elife 10, (2021).

56. Y. Liu, F. M. Ferguson, L. Li, M. Kuljanin, C. E. Mills, K. Subramanian, W. Harshbarger, S. Gondi, J. Wang, P. K. Sorger, J. D. Mancias, N. S. Gray, K. D. Westover, Chemical Biology Toolkit for DCLK1 Reveals Connection to RNA Processing. Cell Chem Biol, (2020).

57. D. Qu, N. Weygant, J. Yao, P. Chandrakesan, W. L. Berry, R. May, K. Pitts, S. Husain, S. Lightfoot, M. Li, T. C. Wang, G. An, C. Clendenin, B. Z. Stanger, C. W. Houchen, Overexpression of DCLK1-AL Increases Tumor Cell Invasion, Drug Resistance, and KRAS Activation and Can Be Targeted to Inhibit Tumorigenesis in Pancreatic Cancer. J Oncol 2019, 6402925 (2019).

58. Y. Ge, H. Liu, Y. Zhang, J. Liu, R. Yan, Z. Xiao, X. Fan, X. Huang, G. An, Inhibition of DCLK1 kinase reverses epithelial-mesenchymal transition and restores T-cell activity in pancreatic ductal adenocarcinoma. Transl Oncol 17, 101317 (2022).

59. S. Thiem, M. F. Eissmann, E. Stuart, J. Elzer, A. Jonas, M. Buchert, M. Ernst, Inducible gene modification in the gastric epithelium of Tff1-CreERT2, Tff2-rtTA, Tff3-luc mice. Genesis 54, 626–635 (2016).

60. K. C. Knower, A. L. Chand, N. Eriksson, K. Takagi, Y. Miki, H. Sasano, J. E. Visvader, G. J. Lindeman, J. W. Funder, P. J. Fuller, E. R. Simpson, W. D. Tilley, P. J. Leedman, J. Graham, G. E. Muscat, C. L. Clarke, C. D. Clyne, Distinct nuclear receptor expression in stroma adjacent to breast tumors. Breast Cancer Res Treat 142, 211–223 (2013).

61. J. Schindelin, I. Arganda-Carreras, E. Frise, V. Kaynig, M. Longair, T. Pietzsch, S. Preibisch, C. Rueden, S. Saalfeld, B. Schmid, J. Y. Tinevez, D. J. White, V. Hartenstein, K. Eliceiri, P. Tomancak, A. Cardona, Fiji: an open-source platform for biological-image analysis. Nat Methods 9, 676–682 (2012).

62. M. J. Goldman, B. Craft, M. Hastie, K. Repecka, F. McDade, A. Kamath, A. Banerjee, Y. Luo, D. Rogers, A. N. Brooks, J. Zhu, D. Haussler, Visualizing and interpreting cancer genomics data via the Xena platform. Nat Biotechnol 38, 675–678 (2020).

63. E. Cerami, J. Gao, U. Dogrusoz, B. E. Gross, S. O. Sumer, B. A. Aksoy, A. Jacobsen, C. J. Byrne, M. L. Heuer, E. Larsson, Y. Antipin, B. Reva, A. P. Goldberg, C. Sander, N. Schultz, The cBio cancer genomics portal: an open platform for exploring multidimensional cancer genomics data. Cancer Discov 2, 401–404 (2012).

64. J. Gao, B. A. Aksoy, U. Dogrusoz, G. Dresdner, B. Gross, S. O. Sumer, Y. Sun, A. Jacobsen, R. Sinha, E. Larsson, E. Cerami, C. Sander, N. Schultz, Integrative analysis of complex cancer genomics and clinical profiles using the cBioPortal. Science signaling 6, pl1 (2013).

65. R. C. Team, “R: A Language and Environment for Statistical Computing,” (R Foundation for Statistical Computing, Vienna, Austria, 2014).

