## Supplementary Figures and Tables for "The kinase activity of the cancer stem cell marker DCLK1 drives gastric cancer progression by reprogramming the stromal tumor landscape"

**This PDF file includes:**

Figs. S1 to S6

Tables S1 to S3

Fig. S1. DCLK1 mRNA expression in gastric cancer cell lines

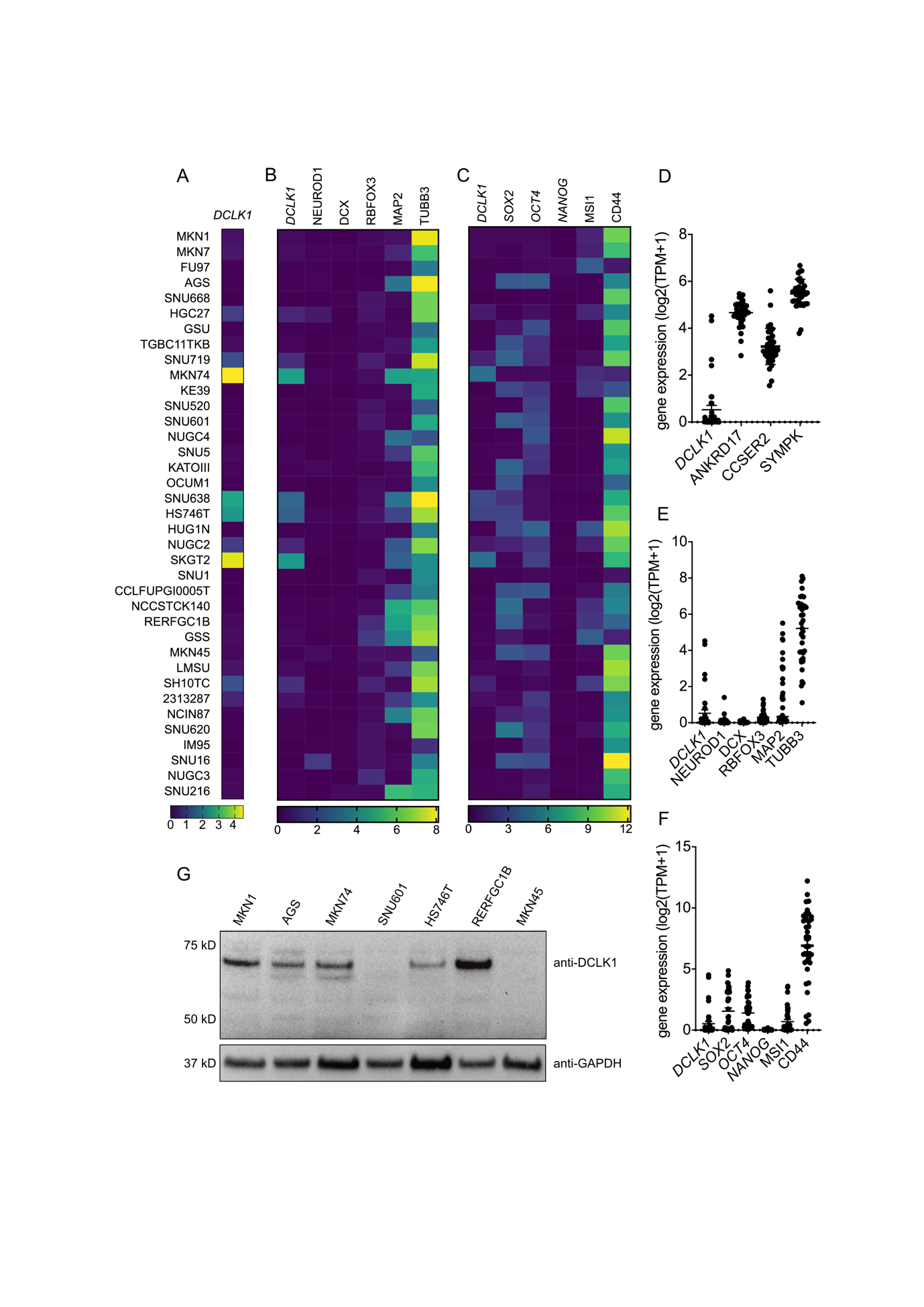

(A) Gene expression heatmap showing DCLK1 (log2(TPM+1)) in 37 gastric cancer cell lines. (B) Gene expression heatmap of DCLK1 and neural markers in 37 gastric cancer cell lines. (C) Gene expression heatmap of DCLK1 and stem cell markers in 37 gastric cancer cell lines. (D) Scatterplot showing transcript levels of DCLK1 and cancer housekeeping genes. (E) Scatterplot showing transcript levels of DCLK1 and neural markers. (F) Scatterplot showing transcript levels of DCLK1 and stem cell markers. (G) Immunoblot of a panel of gastric cancer cell lines for the detection of DCLK1 and GAPDH (loading control).

Fig. S2. DCLK1 promotes growth of gastric cancer xenografts

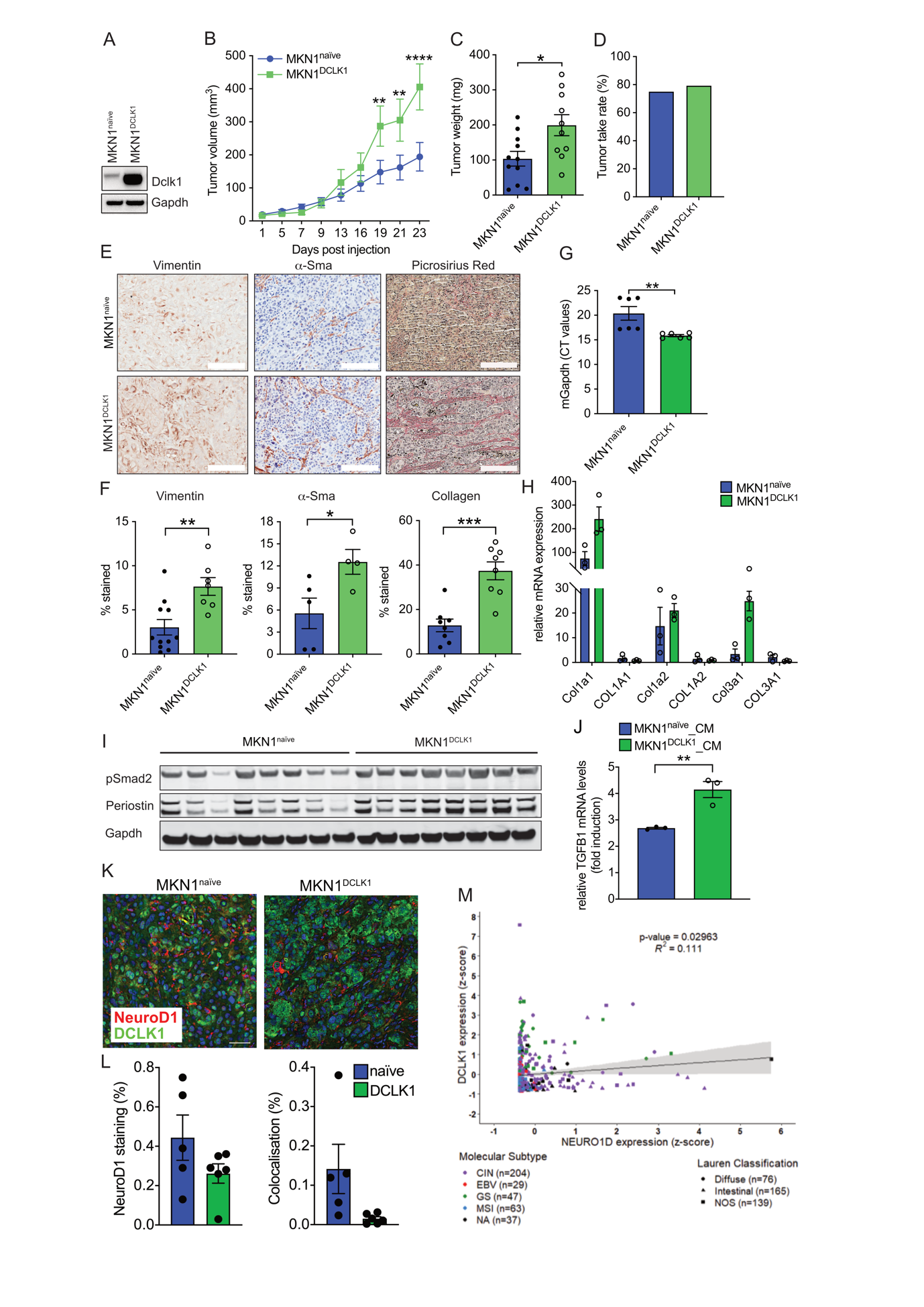

(A) Immunoblot of MKN1^naïve^ and MKN1^DCLK1^ cells lysates for detection of Dclk1 and Gapdh. (B) Xenograft tumor growth of MKN1^naïve^ and MKN1^DCLK1^ cells in Balb/c nude mice. (C) Weight of xenograft tumors at time of harvest. (D) Tumor take rate of MKN1^naïve^ and MKN1^DCLK1^ cells in Balb/c nude mice. (E) Immunohistochemical stains of xenograft tumor tissue using anti-vimentin and α-Sma antibodies and picrosirius red stain to visualize collagen. (F) Quantification of IHC stains. (G) Histogram showing murine Gapdh Ct values. (H) RT-qPCR analysis of xenograft tumors of MKN1^naïve^ and MKN1^DCLK1^ cells using primers specific for the indicated human and murine collagen genes. (I) Immunoblot of MKN1^naïve^ and MKN1^DCLK1^ xenograft tumor lysates for detection of pSmad2, Periostin and Gapdh. (J) RT-qPCR analysis of TGFβ mRNA produced from CAFs stimulated with conditioned medium (CM) prepared from the supernatant of MKN1^naïve^ and MKN1^DCLK1^ cells. (K) Co-immunofluorescence images of tumor xenograft sections stained for DCLK1 and NeuroD1. Nuclei are stained blue using DAPI. (L) Quantification of NeuroD1 staining coverage (%, left) and colocalization with DCLK1 (right) per tumor section in MKN1^naïve^ and MKN1^DCLK1^ tumors. (M) Correlation analysis of DCLK1 and NEUROD1 expression (z-score, log2 transformed) in cancers from the TCGA STAD dataset. Each dot represents a single cancer indicating both its molecular subtype and Lauren classification *P < .05, **P <.01, ***P < .001, ****P <.0001. Scale bar = 300 μm (E), 100 μm (K).

Fig. S3. DCLK1 remodels the tumor stroma in a kinase-dependent manner

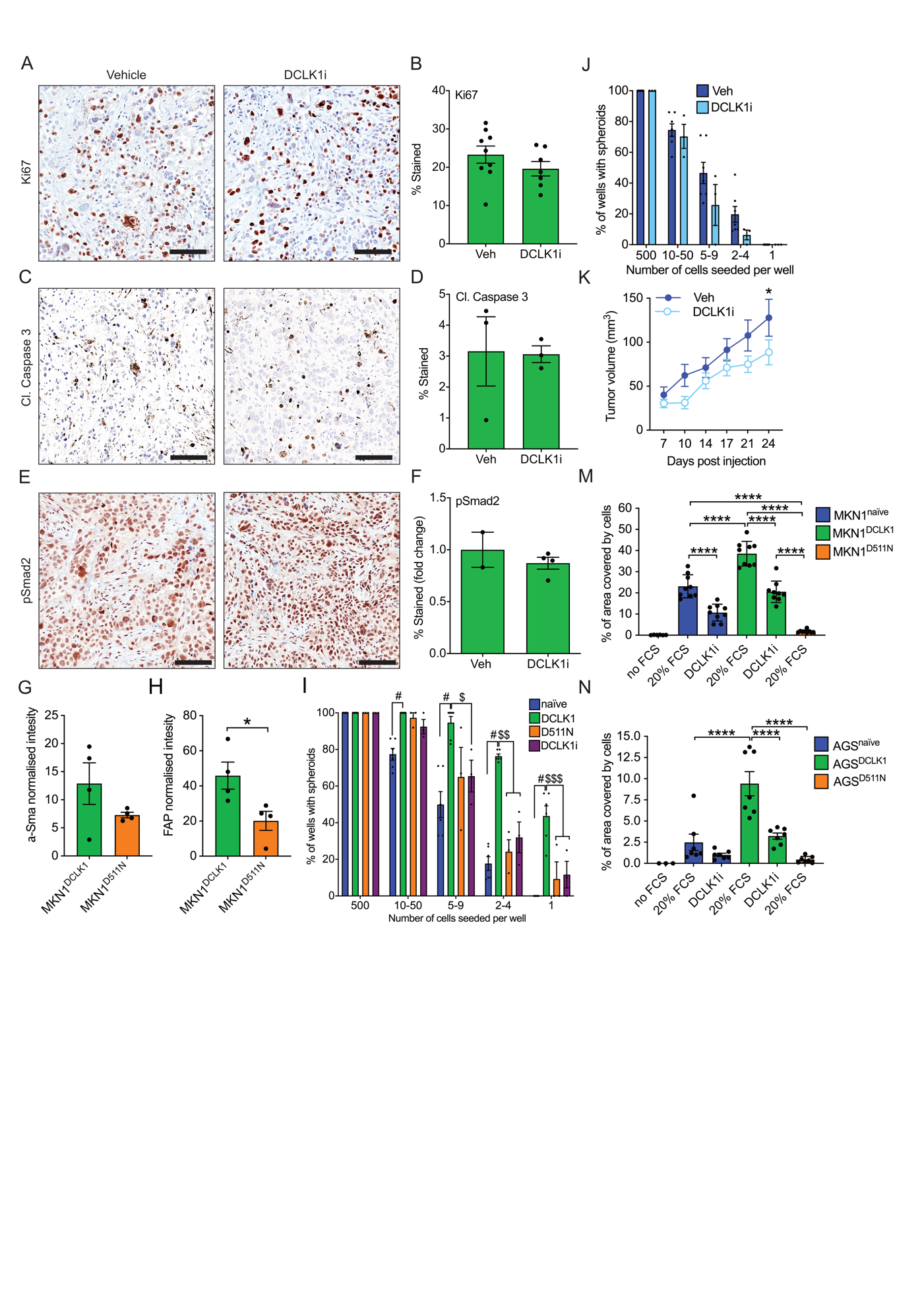

(A) Immunohistochemical stain using anti-Ki67 antibody of xenograft tumors harvested from mice treated with DCLK1 inhibitor (DCLK1i) or vehicle (Veh (B) Quantification of Ki67 IHC stain. (C) Immunohistochemical stain using anti-cleaved Caspase 3 antibody. (D) Quantification of cleaved Caspase 3 IHC stain. (E) Immunohistochemical stain using anti-phospho-SMAD2 antibody. (F) Quantification of phospho-Smad2 IHC stain. (G, H) Quantification of normalised band intensity for α-Sma and Fap. (I) Limiting dilution stem cell assay using MKN1^naïve^, MKN1^D511N^, MKN1^DCLK1^ and MKN1^DCLK1^ treated with DCLK1i. (J) Limiting dilution stem cell assay with MKN1^naïve^ cells treated with DCLK1i or vehicle control. (K) Xenograft tumor growth of MKN1^naïve^ cells in Balb/c nude mice. Mice were treated with DCLK1i or vehicle control. (M) Quantification of cell migration assays using transwell inserts with MKN1^naïve^, MKN1^DCLK1^ and MKN1^D511N^ cells cultured in the presence of 20% FCS or DCLK1i. No FCS is negative control where the FCS is omitted from the lower culture chamber. 20% FCS is the positive control. (N) Quantification of cell migration assays using transwell inserts with AGS^naïve^, AGS^DCLK1^ and AGS^D511N^ cells cultured in the presence of 20% FCS or DCLK1i. Scale bar = 100 μm. *P < .05, ****P <.0001. #, p<0.0001 DCLK1 vs naïve; $, p<0.05 DCLK1 vs DCLK1i; $$, p<0.001 DCLK1 vs D511N and DCLK1i; $$$, p<0.05 DCLK1 vs D511N and DCLK1i.

**Fig. S4.** DCLK1 promotes EMT via CXCL12 chemokine

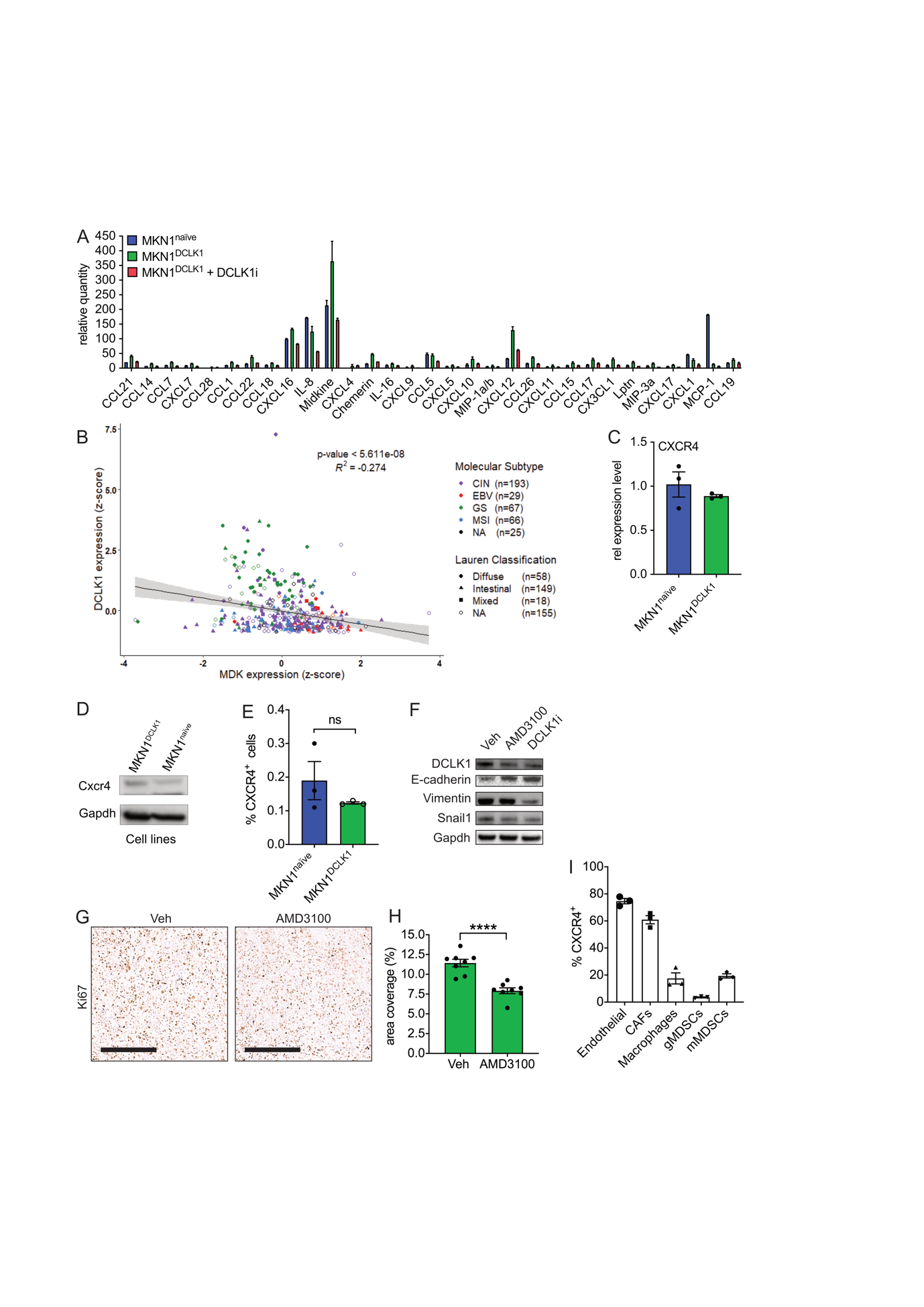

(A) Histogram showing the relative expression levels of chemokines in cell supernatants collected from cultured MKN1^naïve^ and MKN1^DCLK1^ cells and MKN1^DCLK1^ cells grown in the presence of DCLK1 inhibitor for 72 hours (n=2 for MKN1^naïve^ and MKN1^DCLK1^+ DCLK1i; n=4 for MKN1^DCLK1^). (B) Correlation analysis of DCLK1 and Midkine (MDK) expression (z-score, log2 transformed) in gastric cancers from the TCGA STAD dataset. Each dot represents a single cancer indicating both its molecular subtype and Lauren classification. (C) RT-qPCR analysis for human CXCR4 mRNA in MKN1^naïve^ and MKN1^DCLK1^ cells. (D) Immunoblot using anti-CXCR4 antibody on protein lysates of MKN1^naïve^ and MKN1^DCLK1^ cells. (E) Flow cytometric analysis of CXCR4+ cell frequency among MKN1^naïve^ and MKN1^DCLK1^ cancer cells. (F) Immunoblot using antibodies against mesenchymal markers (Vimentin, Snai1) and epithelial marker (E-cadherin) on protein lysates prepared from MKN1^DCLK1^ cells grown for 72 hours in the presence of CXCR4 antagonist (AMD3100), DCLK1 inhibitor (DCLK1i) or vehicle (Veh). (G) Immunohistochemical stain using anti-Ki67 of MKN1^DCLK1^ xenograft tumors harvested from mice treated with AMD3100 or vehicle (Veh). (H) Quantification of Ki67 IHC stains. (I) Flow cytometric analysis of CXCR4+ cell frequency among indicated populations isolated from dissociated MKN1^DCLK1^ xenograft tissue. Scale bar = 300 μm.****P <.0001.

**Fig. S5 (related to Figure 6).** Epithelial and stromal expression of DCLK1 in human diffuse-type GC

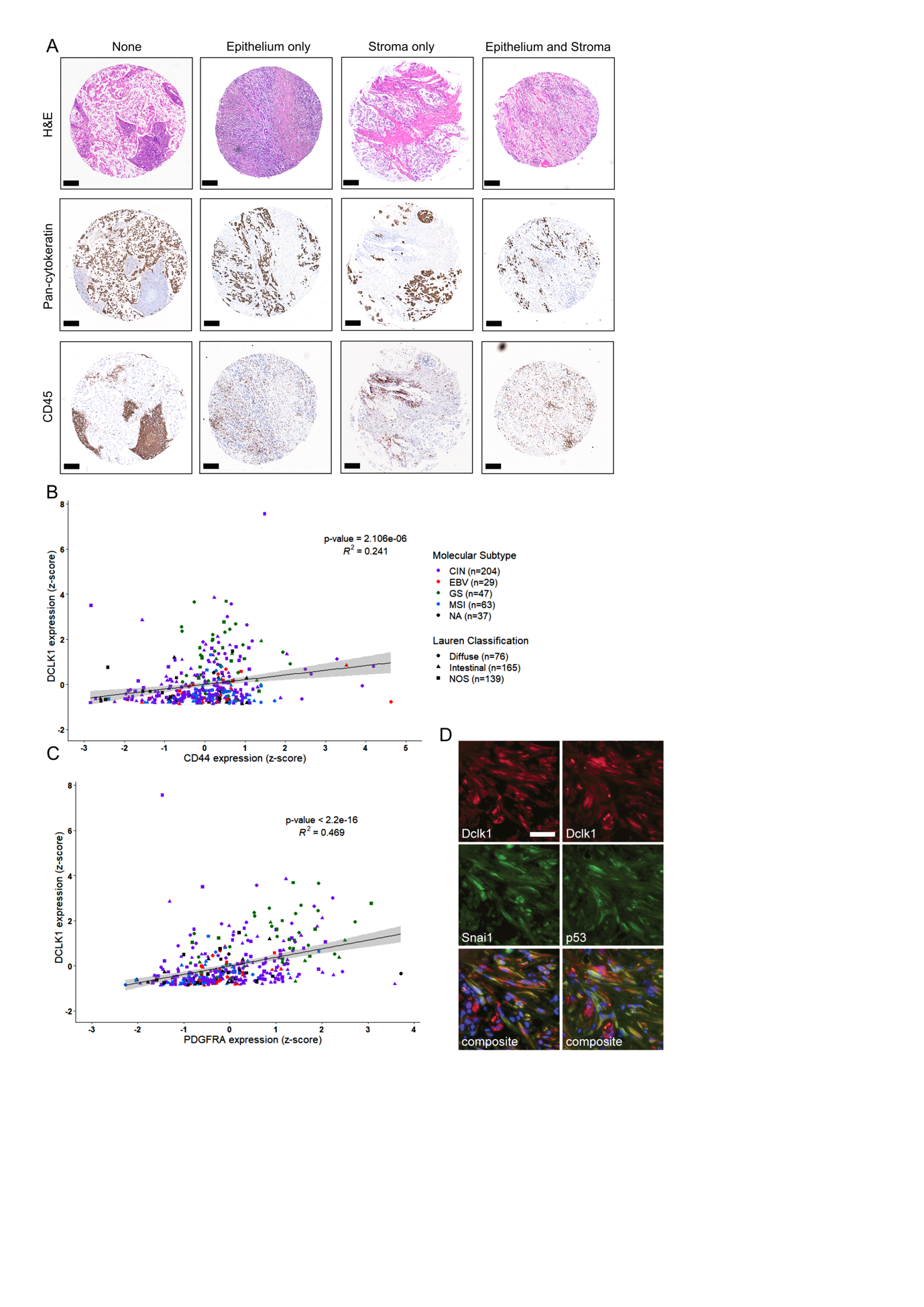

(A) Representative photomicrographs of single core biopsies from a human diffuse-type GC tissue array stained with Hematoxylin and Eosin (H&E, top row), anti-Pan-cytokeratin (middle) or anti-CD45 antibodies (bottom), representing the 4 categories used to assess DCLK1 staining in **Figure 6A**. Shown images belong to same biopsies as used for DCLK1 staining in **Figure 6A**. (B) Correlation analysis of DCLK1 and CD44 expression (z-score, log2 transformed) in gastric cancers from the TCGA STAD dataset. Each dot represents a single cancer indicating both its molecular subtype and Lauren classification. Scale bar = 500 μm. (C) Correlation analysis of DCLK1 and PDGFRA expression (z-score, log2 transformed) in gastric cancers from the TCGA STAD dataset. Each dot represents a single cancer indicating both its molecular subtype and Lauren classification. (D) Co-immunofluorescence staining for Dclk1 and Snai1 (left panel) and Dclk1 and p53 (right panel) on sections of a murine invasive gastric cancer grown in *Tff1CreERT2;Kras^G12D^;Pi3kca^H1047R^;p53^R172H^* mice. Nuclei are stained blue using DAPI. Scale bar = 100 μm.

**Fig. S6 (related to Figure 7).** Spearman correlation matrix of DCLK1 with functional gene signatures using the TCGA-STAD dataset

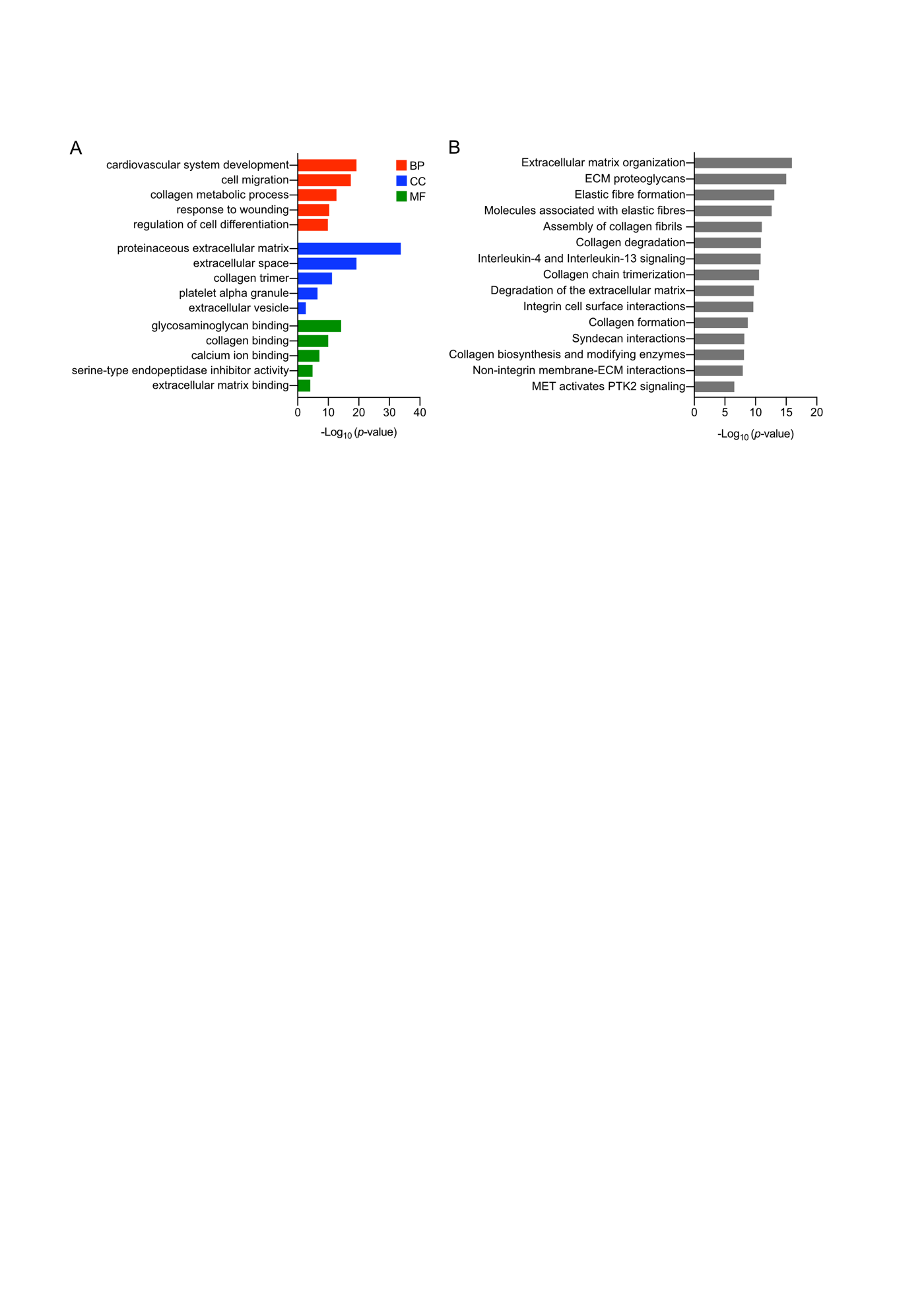

(A) GO-term analysis of the genes within the boxed area in Figure 7A. Shown are the -Log10 p-values of the top 5 GO-terms with respect to biological processes (BP), cellular components (CC) and molecular functions (MF). (B) Reactome analysis of the genes corresponding to the boxed area in Figure 7A. Shown are the 15 most significantly enriched pathways.

Table S1.

| **Gene** | **Forward 5’-3’** | **Reverse 3’-5’** |
| --- | --- | --- |
| Gapdh_m.m | aagagggatgctgccctta | ttttgtctacgggacgagga |
| Hprt_m.m | tcctcctcagaccgctttt | cctggttcatcatcgctaatc |
| HPRT_h.s | gaccagtcaacaggggacat | gtgtcaattatatcttccacaatcaag |
| CXCR4_h.s | ggatataatgaagtcactatgggaaaa | agtagtgggctaagggcaca |
| DCLK1_h.s | cccaagccgaatagcaca | ccttatcaagagcggtggtt |
| Dclk1_m.m | ttcaacacaggccccaag | tatcaagagcggtggttgc |
| COL1A1_h.s | ctggacctaaaggtgctgct | gctccagcctctccatctt |
| Col1a1_m.m | agacatgttcagctttgtggac | gcagctgacttcagggatg |
| COL1A2_h.s | tctggagaggctggtactgc | ccagggagacccagaatacc |
| Col1a2_m.m | caagcatgtctggttaggagag | aggacaccccttctacgttgt |
| COL3A1_h.s | agggaaagatggcccaag | ggcaccaccttcaccctta |
| Col3a1_m.m | tcccctggaatctgtgaatc | tgagtcgaattggggagaat |
| TGFB1_h.s | cgctgcccatcgtgtacta | cgcacgatcatgttggac |
| SNAI1_h.s | gcgagctgcaggactctaat | cggtggggttgaggatct |
| SNAI2_h.s | tggttgcttcaaggacacat | gcaaatgctctgttgcagtg |

RT-qPCR primers used in this study.

Table S2.

| **Antibody/Kit** | **Supplier** | **Cat. No** | **Species** | **Used for / Dilution** |
| --- | --- | --- | --- | --- |
| Ki67 | Thermo Scientific | MA5-14520 | Rabbit | IHC / 1:100 |
| α-Smooth Muscle Actin | Abcam | ab5694 | Rabbit | IHC / 1:100 |
| Vimentin | Cell Signaling | CST3932S | Rabbit | IHC / 1:100 |
| Dclk1 | Abcam | ab106635 | Rabbit | IHC/ 1 :1000 |
| ApopTag Plus Peroxidase *In Situ* | Millipore | S7101 | N/A | IHC / 1:10 |
| Anti-human mitochondria | Sigma-Aldrich | MAB1273 | Mouse | IHC / 1:500 |
| Lysyl oxidase (LOX) | Santa Cruz | sc-373995 | Mouse | IHC / 1:100 |
| Phospho-Smad2 (pSmad2) | Abcam | ab188334 | Rabbit | IHC / 1:200 |
| Cleaved Caspase 3 | Cell Signaling | CST96612 | Rabbit | IHC / 1:100 |
| Pan-Cytokeratin | Abcam | ab27988 | Mouse | IHC / 1:40 |
| p53 | Santa Cruz | sc-98 | Mouse | IF / 1:150 |
| Snai1 | Santa Cruz | sc-393172 | Mouse | IF / 1:250 |
| Dclk1 | Abcam | ab31704 | Rabbit | IF / 1:200 |
| NeuroD1 | Santa Cruz | sc-46684 | Mouse | IF / 1:200 |
| Dclk1 | Abnova | H00009201 | Mouse | WB / 1:1000 |
| Snai1 | Abclonal | A5544 | Rabbit | WB / 1:1000 |
| Cxcl12 | Abcam | ab9797 | Rabbit | WB / 1:1000 |
| Cxcr4 | Invitrogen | 2B11 | Mouse | WB / 1:300 |
| Gapdh | Sigma-Aldrich | G9545 | Rabbit | WB / 1:2000 |
| α-tubulin | Abcam | ab7291 | Mouse | WB / 1:1000 |
| pAkt | Cell Signaling | CST4058 | Rabbit | WB / 1:500 |
| pErk | Cell Signaling | CST9101 | Rabbit | WB / 1:1000 |
| pSmad2 | Abclonal | AP0548 | Rabbit | WB / 1:1000 |
| p53 | Thermo Scientific | PA6240 | Rabbit | WB / 1:1000 |
| Fap | Abcam | ab53066 | Rabbit | WB / 1:500 |
| Periostin | Santa Cruz | sc-398631 | Mouse | WB / 1:1000 |
| E-cadherin | Abcam | ab15148 | Rabbit | WB / 1:500 |
| Vimentin | Abcam | ab92547 | Rabbit | WB / 1:1000 |

Antibodies used for IHC, IF and WB.

Table S3.

| **Antibody** | **Supplier** | **Clone** | **Species** | **Dilution** |
| --- | --- | --- | --- | --- |
| CD45-PerCP/Cy5.5 | Biolegend | 2D1 | Mouse | 1:200 |
| EPCAM-FITC | Biolegend | CO17-1A | Mouse | 1:200 |
| CD31-PE/CY7 | Biolegend | 390 | Rat | 1:200 |
| PDPN-APC/CY7 | Biolegend | 8.1.1 | Hamster | 1:200 |
| PDGFRA-PE | Biolegend | 16A1 | Mouse | 1:200 |
| LY6C-Efluor 450 | Invitrogen | RB6-8C5 | Rat | 1:200 |
| LY6G-PerCp/Cy5.5 | Biolegend | 1A8 | Rat | 1:200 |
| CD11B-PE | Biolegend | M1/70 | Rat | 1:200 |
| F4/80-FITC | Biolegend | BM8 | Rat | 1:200 |
| CXCR4-APC | Biolegend | L276F12 | Rat | 1:100 |
| SYTOX Blue | Thermo Fisher | N/A | N/A | 1:250 |

Antibodies used for flow cytometry.
